# Female-enriched *Eggerthella lenta* drives neuroinflammation and IFN-γ via host receptor TLR2

**DOI:** 10.64898/2026.03.16.711194

**Authors:** Rachel R. Rock, Margaret Alexander, Cecilia Noecker, Kai R. Trepka, Vaibhav Upadhyay, Edwin F. Ortega, Lorenzo Ramirez, Lena Siewert, Christine A. Olson, Taylor Halsey, Anne-Katrin Pröbstel, Sergio E. Baranzini, Peter J. Turnbaugh

**Affiliations:** Department of Microbiology & Immunology, University of California, San Francisco, San Francisco, CA USA; Department of Medicine, University of California, San Francisco, San Francisco, CA USA; Center for Neurology & Clinic for Neuroimmunology and Neuromuscular Diseases, University Hospital and University of Bonn, Bonn, Germany; Departments of Neurology, Biomedicine and Clinical Research, and Research Center for Clinical Neuroimmunology and Neuroscience Basel (RC2NB), University Hospital Basel and University of Basel, Basel, Switzerland; Department of Neurology, Weill Institute for Neurosciences, University of California San Francisco, San Francisco, CA USA; Biohub, San Francisco, CA USA

**Keywords:** multiple sclerosis, gut-brain axis, sex as a biological variable, human gut microbiome, meta-analysis, *Eggerthella lenta*, autoimmune disease, T helper 1 cells, T helper 17 cells, Toll-like receptor 2

## Abstract

Women are at increased risk of autoimmune diseases, including multiple sclerosis (MS); however, the degree to which sex differences in the gut microbiota impact autoimmunity remains largely unexplored. Our 27-cohort meta-analysis revealed 60 sex-associated gut bacterial species. Leveraging an independent clinical cohort, we demonstrate that female-enriched species significantly associate with MS status and clinical disability (EDSS). Top female-enriched species *Eggerthella lenta* drove disease in the experimental autoimmune encephalomyelitis (EAE) MS model, consistent with brain and gut lamina propria T cell infiltration and MS-associated T helper (Th) signatures. *E. lenta* induced intestinal Th1 and Th17 in healthy mice, independent of bacterial viability. Mechanistically, we demonstrate that TLR2 directly drives *E. lenta*-induced IFN-γ production in Th cells and is necessary for exacerbation of EAE. Together, we identify a causal host-microbe axis contributing to sex differences in autoimmunity and provide a framework for evaluating sex as a biological variable in human microbiome research.

**HIGHLIGHTS:**

- 27-cohort meta-analysis identifies a robust sex-signature in human gut microbiota.
- Female-enriched species are associated with MS risk and severity.
- Female-enriched *Eggerthella lenta* exacerbates the EAE model.
- *E. lenta* impacts neuroinflammation via toll-like receptor 2.

## INTRODUCTION

Most patients with autoimmune disease are women, with female sex approximately doubling incidence^1^. This is also true for multiple sclerosis (MS), a central nervous system (CNS) autoimmune disease where the immune system attacks the critical brain protein myelin. MS is at least twice as prevalent in women than in men^2,3^ and this sex bias continues to increase with age^3^. While there has been extensive prior literature exploring the role of environmental factors like vitamins^2,3^, obesity^4^, and smoking^2^ in MS, none of these factors explain the observed sex bias in disease risk and severity^2,3^. Instead, we hypothesized that the trillions of microorganisms found within the gastrointestinal tract (the gut microbiota) could play an underappreciated role in contributing to sex bias in MS and potentially other autoimmune diseases.

Studies in mouse models and people living with MS (PwMS) have implicated the gut microbiota in MS pathogenesis^5–8^. The widely-studied experimental autoimmune encephalomyelitis (EAE) mouse model of neuroinflammation is primarily driven by T helper 1 (Th1)^9–11^ and T helper 17 (Th17) cells^11,12^, both of which depend on the gut microbiota^13^. Germ-free (GF) mice, which fully lack microbiota, are protected from EAE and conventionalization (re-colonization of formerly GF mice) is sufficient to restore disease^5,6^. Transplantation of ileal or stool microbiota from PwMS significantly exacerbates EAE phenotypes relative to recipients of healthy control samples^7,8,14^. A highly limited set of bacteria enriched in PwMS are known to exacerbate EAE, including *Tyzzerella nexilis*^15^ and, in certain contexts, *Akkermansia muciniphila*^16^. Other members of the gut microbiota are protective, including *Prevotella histolytica*^17^ and ketogenic diet-associated *Lactobacillus*^18^. Taken together, these results support a model wherein the balance between specific gut bacterial species contributes to the risk and progression of disease.

However, these prior studies have not adequately considered the role host sex plays in shaping overall gut microbial community structure and/or the abundance of specific immunomodulatory gut bacterial species. Here, we address this critical knowledge gap. Leveraging high-quality, multi-cohort metagenomic data, we reveal a consistent signature of biological sex in the human gut microbiota. We then focus on *E. lenta* as a *proof-of-concept* for the causal role of a female-enriched gut bacterial species in disease. We report the first instance of a human gut Actinomycetota that drives EAE, adding *E. lenta* to the growing list of gut bacterial species relevant to MS and ultimately providing a new framework to explore the microbial etiology of sex disparities in disease risk, progression, and treatment outcomes.

## RESULTS

### Identification of sex-associated bacterial species enriched in autoimmune disease

While sex is frequently recorded as a variable in individual microbiota profiling studies, we sought to identify gut microbial species robustly associated with sex across multiple independent studies and cohorts. We leveraged the <u>c</u>urated <u>M</u>etagenomics <u>D</u>ata resource^19^ (cMD) to re-analyze metagenomic data from 3,979 subjects across 27 independent cohorts (**Table S1**). Sample inclusion criteria included no documented antibiotic use, complete metadata for model covariates, and both high sequencing coverage and quality (see STAR Methods). After adjusting for confounding variables, we identified 91 differentially abundant bacterial species between male and female subjects (**Figure 1A** and **Data S1**) and 149 bacterial species and one archaeal species with a significant difference in prevalence (**Figure S1A** and **Data S2**). Combined, we identified 166 distinct species associated with sex when considering abundance and/or prevalence (**Figure 1B**), including 101 female-enriched and 65 male-enriched species (**Table S2**).

**Figure 1.**
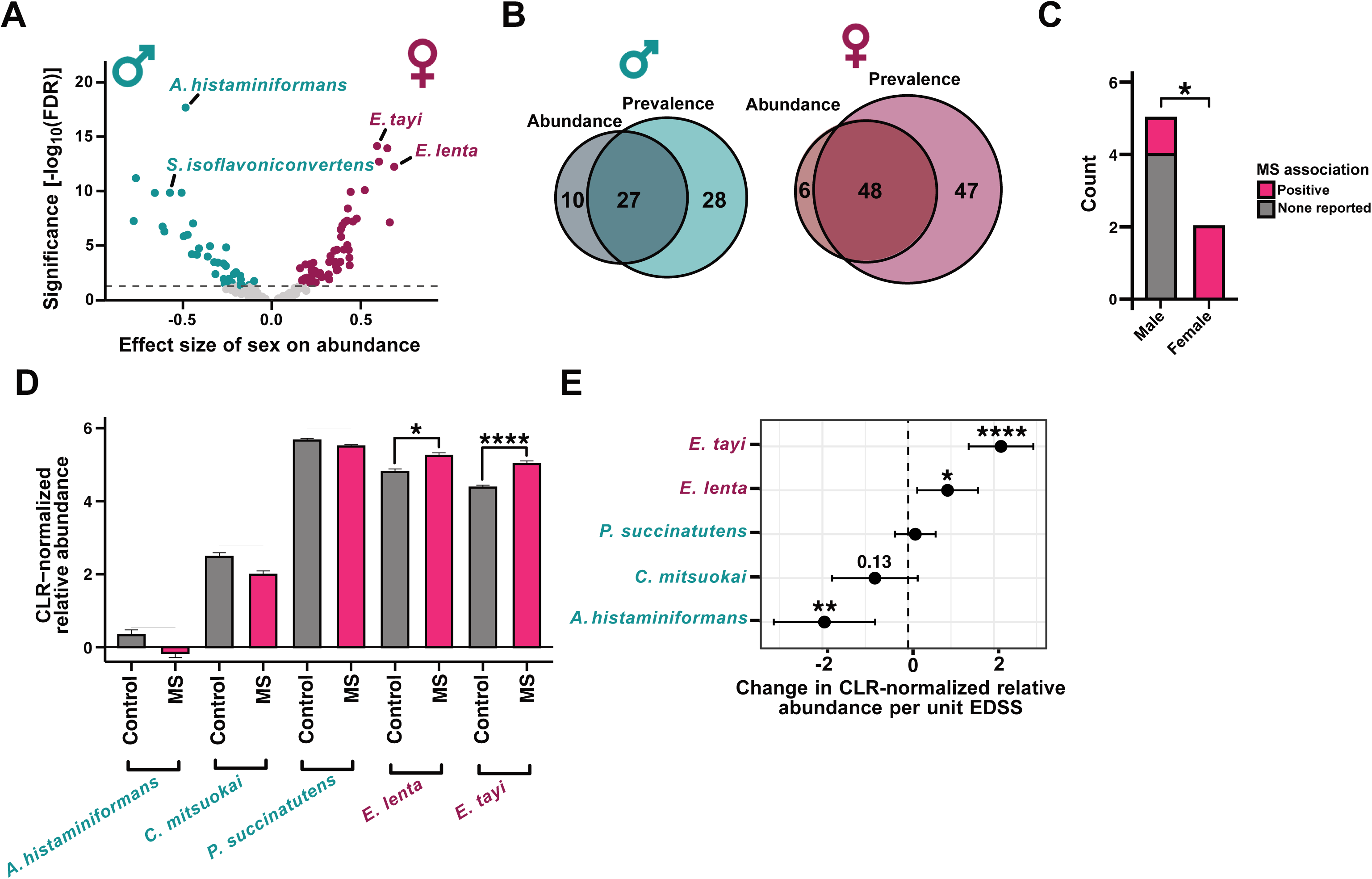
**Multiple human gut bacterial species show sex-specific enrichment and are associated with multiple sclerosis.** (A and B), A published metagenomic data repository representing metagenomic samples from 14,859 individuals was filtered to include only high-quality stool samples (n=4,681 samples) and re-analyzed to identify bacterial species associated with sex. MaAsLin2 linear models were fit with sex (male or female), age category, BMI category, continent, health category, median read length category, sequencing depth category, and DNA extraction kit category as fixed effects and study as a random effect (CLR-normalized abundance ∼ sex + age category + BMI category + continent + health category + median read length category + sequencing depth category + DNA extraction kit category + (1|study), see also STAR Methods). (A) We identified 91 differentially abundant bacterial species enriched in female (mauve) or male (teal) individuals (FDR < 0.05). MaAsLin2 linear models were fit using CLR-normalized abundance with sex, age, BMI, continent, health category, and sequencing metrics as fixed effects and study as a random effect. (B) Intersection of results from (A) with a model of species prevalence. Prevalence was determined via FDR-adjusted logistic regression models (see also Figure S1A). (C) Analysis of the MetaBiom database revealed that female-enriched bacteria (see Figure S1D) are significantly more likely to be MS-associated (counts displayed; see also Table S3). (D and E) Relative abundance of sex-associated species in the International Multiple Sclerosis and Microbiome Consortium (iMSMS) dataset (n=576 cases, n=576 controls). Female-enriched species are (D) higher in MS patients and (E) positively associated with disease severity (expanded disability status score [EDSS]). p values are likelihood ratio tests (D) or sex- and treatment-adjusted linear models (D, E).Mean ± model SEM (D) or mean ± 95% model CI (E) are displayed. See also Figure S1 and Tables S1–S3. *p < 0.05, **p < 0.01, ***p < 0.001, ****p < 0.0001. Full species names: *Allisonella histaminiformans, Catenibacterium mitsuokai, Eggerthella lenta, Eisenbergiella [Dialister] tayi, and Phascolarctobacterium succinatutens*.

We further narrowed down this list by applying more stringent filtering criteria, focusing on the microbial species in the top quintiles for effect size and bottom quintiles for significance (**Figs. S1B-D**). This analysis revealed a total of 60 high-confidence sex-associated microbial species, including 34 female-enriched and 26 male-enriched species (**Figure S1D** and **Table S2**). Notably, we identified 5 male-enriched and 2 female-enriched species (*Eisenbergiella tayi* and *Eggerthella lenta*) that were consistently identified as top hits by both abundance and prevalence.

Both female-enriched bacterial species were enriched in PwMS and positively associated with disease severity. We analyzed the overall association between each of the top 7 sex-associated bacterial species and MS within the MetaBiom database^20^. The female-enriched *E. tayi* and *E. lenta* were both enriched in PwMS, whereas 4/5 male-enriched bacterial species were not associated with MS (**Figure 1C** and **Table S3**). To further validate these observations, we compared the abundance of each species in PwMS and healthy controls from the International Multiple Sclerosis and Microbiome Study (iMSMS) cohort^21^. Consistent with the MetaBiom-based analysis, both female-enriched bacterial species were significantly enriched in PwMS relative to controls after adjusting for sex and treatment status (**Figure 1D**). None of the male-enriched species detected in iMSMS were significantly different between PwMS and controls (**Figure 1D**). Furthermore, both female-enriched species were positively associated with disease severity after adjusting for sex and treatment status (**Figure 1E**). Notably, the male-enriched species *Allisonella histaminiformans* was negatively associated with disease severity (**Figure 1E**), suggesting a potential protective effect. We validated these findings using an alternative criterion for the top sex-associated bacteria (**Figure S2**), confirming that female-enriched gut bacterial species exhibit a striking and positive association with disease status and severity in human subjects.

### The female-enriched gut bacterium E. lenta exacerbates neuroinflammation in mice

Given the *E. lenta* enrichment we observed in MS, we tested the impact of the *E. lenta* type strain (DSM2243) on EAE phenotypes in mixed-sex C57BL/6J mice dosed with the MOG_35-55_ antigen followed by two doses of pertussis toxin^22^ (n=32 mice/group). Live *E. lenta* was grown in a chemically-defined and myelin-free media^23^ for every-other-day oral gavage starting two weeks prior to EAE induction (**Figure 2A**). As expected, *E. lenta* was detectable only in endpoint stool samples from the *E. lenta*-gavaged mice (**Figure 2B**). Disease incidence was comparable between groups (**Figure 2C**), indicating that *E. lenta* is not required for EAE induction in the context of a complex microbiota. However, *E. lenta* administration led to significantly increased disease scores over time (**Figure 2D**), resulting in a significantly higher maximum disease score of affected mice (**Figure 2E**) and more severe outcomes by score distribution (**Figure 2F**). While male mice trended towards more severe phenotypes, no significant effect of sex on disease course was observed (**Figure S3**). Colonization of GF mice with *E. lenta* DSM2243 was sufficient to elicit early manifestations of EAE (**Figure S4**), with a significantly higher incidence rate and disease scores relative to GF controls. However, disease phenotypes were very mild overall, consistent with prior data showing less severe EAE in gnotobiotic mice^6,8^.

**Figure 2.**
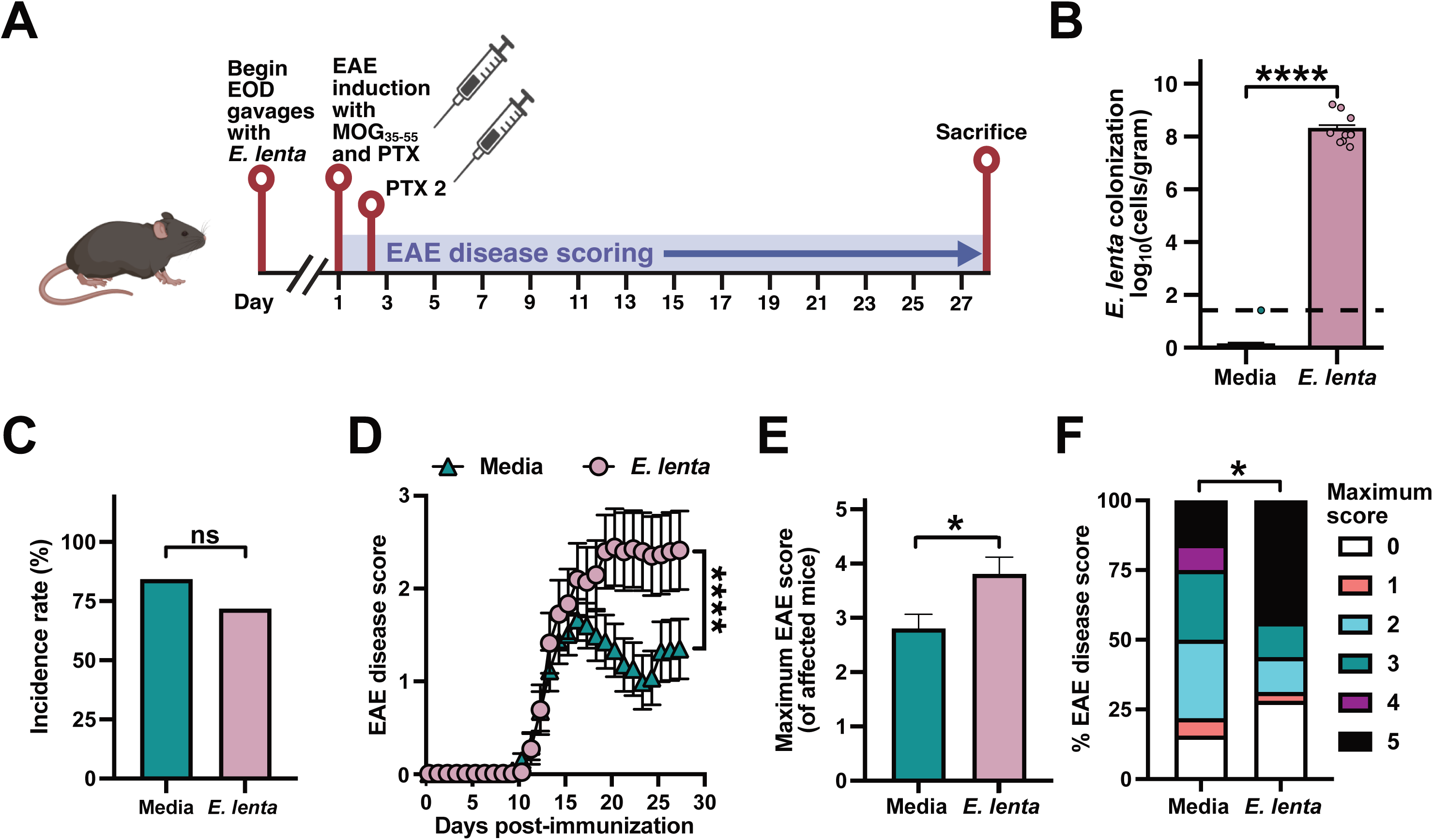
***E. lenta* exacerbates EAE in CONV-R mice.** (A) Mixed-sex, pair-housed adult C57BL/6J CONV-R mice were gavaged every-other-day [EOD] with media or *E. lenta* DSM2243 (n=32 mice/group total from 2 independent experiments). (B) *E. lenta* levels (log_10_ cells/gram) in endpoint cecal contents measured by qPCR (n=10–15 mice/group). Dashed line indicates the limit of detection; p values are Wilcoxon rank sum tests. (C–F) EAE phenotypes were tracked for: (C) incidence, (D) disease score, (E) maximum severity of mice that develop disease, and (F) proportion of mice at peak disease scores. p values are likelihood ratio tests of a global generalized linear model (GLM) with binomial error and logit link (C), repeated measures mixed-effects models with mouse as a random effect (D), Wilcoxon rank sum tests (E), or Fisher’s exact tests (F). Values are mean +SEM (B, E), percentage (C, F), or mean ± SEM (D). *p < 0.05, ****p < 0.0001. See also Figure S3.

Previously, we demonstrated that the cardiac glycoside reductase (*cgr*) operon is necessary for *E. lenta* to activate colonic Th17 cells^24,25^. Given the well-established role of Th17 cells in EAE^5,6^, we hypothesized that the *cgr* operon may also be required for *E. lenta* to exacerbate EAE. We replicated the EAE model with repeated oral gavage of wild-type (wt) *E. lenta*, Δ*cgr E. lenta*, and media controls, ending the experiment at peak disease to enable flow cytometry-based quantification of immune cells (**Figure 3A**). While *E. lenta* was detectable in mice colonized with both the wt and Δ*cgr* strain, but not media controls (**Figure 3B**), *cgr2* was only detected in the ceca of mice colonized with wt *E. lenta* (**Figure 3C**). Neither wt nor Δ*cgr E. lenta* increased incidence rate (**Figure 3D)**, but both strains robustly drove EAE disease score (**Figure 3E**), with a higher maximum disease score of affected mice (**Figure 3F**) and more severe outcomes by score distribution (**Figure 3G**) relative to media controls. This result indicates that the impact of *E. lenta* on EAE is independent of the *cgr* operon, prompting us to consider other aspects of host immunity that are influenced by *E. lenta* in the context of disease.

**Figure 3.**
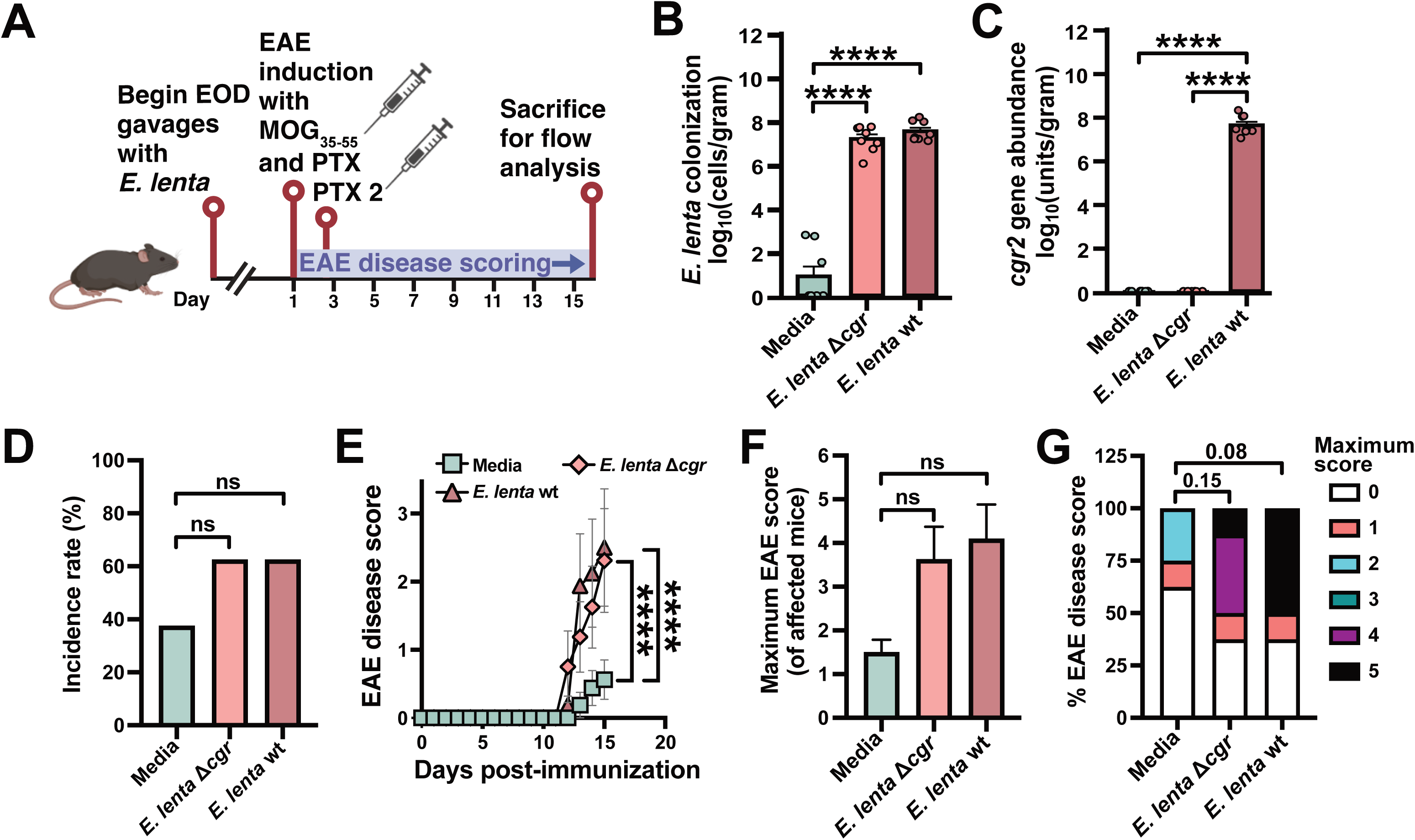
***E. lenta*-induced EAE disease is independent of the cgr operon.** (A) Male C57BL/6J mice were gavaged with media, Δ*cgr E. lenta* DSM 2243, or wt *E. lenta* DSM 2243(n = 8 mice/group). (B and C) Endpoint cecal levels of (B) E. lenta and (C) the *cgr2* gene cluster measured by qPCR (n = 7-8 mice/group). p values are Wilcoxon rank sum tests. (D-G) EAE clinical phenotypes: (D) incidence, (E) disease score, (F) maximum disease severity, and (G) peak score proportions. p values are likelihood ratio tests of a global GLM (D), repeated measures mixed-effects models (E), Wilcoxon rank sum tests (F), or Fisher’s exact tests (G). Values are mean + SEM (B, C, F), percentage (D), or mean ± SEM (E). ****p < 0.0001. See also Figure S5.

We utilized flow cytometry to assess the extent of immune infiltration in the colon lamina propria and brain at peak disease. Notably, both wt and Δ*cgr E. lenta* significantly increased T cell levels in both tissues (**Figs. S5A-D**). Th17 master transcription factor RORγt trended higher in both tissues in response to *E. lenta* (**Figs. S5E-H**) and the Th17 signature cytokine IL-17A was significantly increased in the colonic lamina propria but not in the brain (**Figs. S5I-L**). Notably, the Th1 signature cytokine IFN-γ was markedly and significantly increased in colonic lamina propria and brain tissue for both wt and Δ*cgr E. lenta*-treated mice (**Figs. S5M-P**).

These differences in host immunity during peak disease prompted us to more comprehensively assess the impact of *E. lenta* on intestinal immunity during homeostasis. To assess Th1 response, we re-analyzed bulk RNA sequencing (RNA-seq) data from GF and *E. lenta* DSM2243 monocolonized mouse ileal cells expressing helper T cell marker CD4^25^. Consistent with our original analyses^25^, “Th17 cell differentiation” was the most significantly enriched pathway when comparing the genes differentially expressed between *E. lenta* monocolonized and GF controls (**Figure 4A**). However, the combined pathway for “Th1 and Th2 cell differentiation” was also significantly enriched, including multiple Th1 hallmark genes (**Figs. 4A-C**). We confirmed these findings using an alternative pathway enrichment tool (**Figure S6**). We also validated the impact of *E. lenta* on Th1 response using flow cytometry. *E. lenta* was sufficient to induce IFN-γ, a key Th1 effector, in both lamina propria from ilea (**Figs. 4D,E**) and colons (**Figure 4F,G**) of gnotobiotic mice. We also observed a significant increase in both CD4^+^IFN-γ^+^ and CD4^-^IFN-γ^+^ among live, singlet T cells in the ileum (**Figure S7A-C**) and significantly increased IFN-γ fluorescence within both the CD4^+^ and CD4^-^ T cell populations (**Figure S7D-G**) for both tissues. Taken together, these results support a model in which *E. lenta* predisposes mice to more severe EAE by increasing the level of intestinal Th1 cells, which can then exacerbate neuroinflammation upon disease induction.

**Figure 4.**
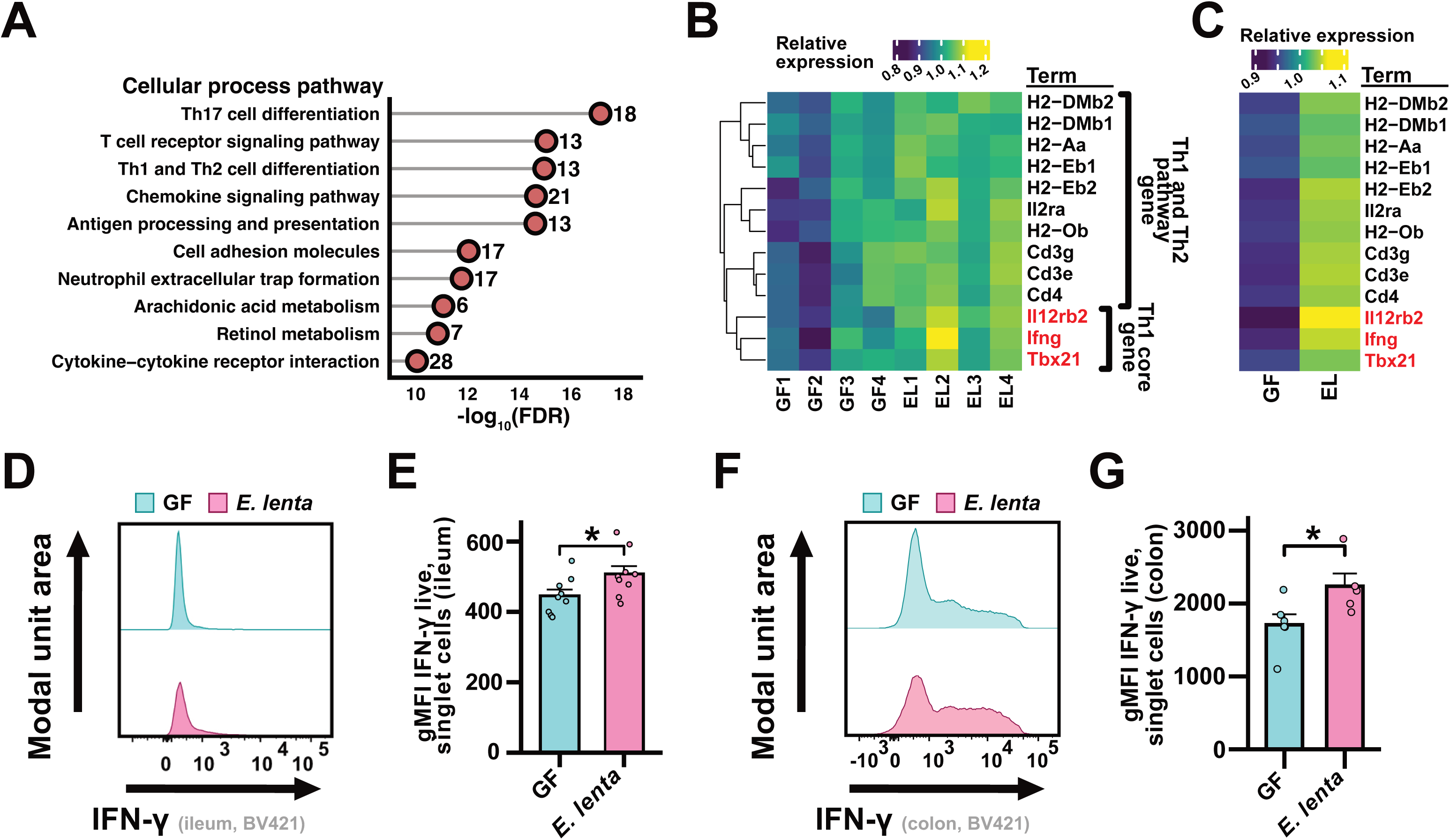
***E. lenta* is sufficient to induce interferon-gamma and Th1-type immune response in germ-free mice.** (A) Pathway enrichment analysis of published RNA-seq data from ileal CD4^+^ T cells (n = 4 mice/group). DEGs were identified via DESeq2 (|FC| > 1.5, Wald test p < 0.05) and analyzed for cellular process terms; p values are Fisher’s exact tests (FDR shown; see also Table S10). (B and C) Relative expression (variance stabilized transformation [VST] values) of differentially expressed genes (DEGs) from the Th1 and Th2 cell differentiation pathway displayed by (B) sample and (C) group average. Core Th1 genes are highlighted in red. (D-G) Flow cytometry analysis of the (D, E) ileal and (F, G) colonic lamina propria from monocolonized and GF mice (n = 5-9 mice/group). (D, F) Representative histograms and (E, G) gMFI quantification of IFN-γ. p values are Wilcoxon rank sum tests. Mean + SEM is displayed; each point represents an individual mouse. *p < 0.05. See also Figures S6 and S7 and Tables S9 and S10.

### Induction of IFN-γ by E. lenta cell components requires TLR2

To better understand the mechanism through which *E. lenta* drives neuroinflammation we tested the impact of *E. lenta* DSM2243 in the SJL/J model of EAE. This model utilizes the proteolipid protein fragment PLP_139-151_ to induce EAE in the SJL/J background (**Figure S8A**). As expected, *E. lenta* was detectable in the cecum at our experimental endpoint (**Figure S8B**). Consistent with MOG_35-55_ in the C57BL/6J background (**Figure 2C**), EAE incidence was comparable between groups (**Figure S8C**). However, in contrast to the MOG_35-55_ C57BL/6J model, disease severity was not significantly different between groups (**Figs. S8D-F**). Stratifying by sex revealed a significant decrease in disease severity in *E. lenta*-treated females (**Figure S9**). Taken together, these results indicate that the ability of *E. lenta* to exacerbate EAE is specific to the MOG_35-55_ C57BL/6J model.

We initially hypothesized that the failure of *E. lenta* to drive disease in this alternative model could reflect antigen mimicry, specifically that *E. lenta* sensitizes T cells to MOG_35-55_ but not PLP_139-151_. We leveraged pattern-matching code (see STAR Methods) to search for immune epitopes that could cross-react with MOG_35-55_ within the *E. lenta* DSM2243 genome. Our search revealed putative mimicry motifs in 13 distinct proteins (**Table S4**). However, an *in vitro* assay for MOG_35-55_ mimicry showed no difference in response to *E. lenta* cells or cell products (**Figure S10A,B,D,E**) or either of two representative *E. lenta* proteins (**Figure S10C,F**) despite robust responsiveness to the positive control.

An alternative hypothesis for the differences we observed in the two tested EAE models is that the distinct genetic background of the mice is responsible. Notably, SJL mice carry a well-characterized amino acid substitution in TLR2 that alters ligand responsiveness and downstream signaling, which has been shown to contribute to their divergent inflammatory and clinical EAE disease phenotypes relative to the C57BL/6J strain^26,27^. To test this hypothesis, we first harvested CD4^+^ T cells from C57BL/6J mice and cultured them under Th1 polarizing conditions in the presence or absence of heat-killed *E. lenta* DSM2243 cells. In this assay, *E. lenta* cells significantly increased IFN-γ (**Figure 5A**). This enhanced skewing effect is consistent with the observation that in healthy human adults, *E. lenta* relative abundance in the gut correlates with heightened IFN-γ response following *M. tuberculosis* stimulation^28^. To assess the generalizability and specificity of this *in vitro* effect, we tested a panel of 8 strains from the *Actinomycetota* phylum, including 4 strains of *E. lenta*. The level of IFN-γ was significantly higher in response to *E. lenta* than strains from other genera, with *E. lenta* DSM2243 exhibiting the highest level of induction (**Figure 5B**). Finally, we directly tested the impact of TLR2 in this assay using T cells harvested from *tlr2*-deficient (*tlr2^-/-^*) C57BL/6J mice and FSL-1, a TLR2 agonist that acts through potent activation of the TLR2/TLR6 receptor complex. Consistent with our prior data, both heat-killed *E. lenta* and FSL-1 significantly increased IFN-γ in wild-type cells (**Figure 5C**). However, IFN-γ levels were significantly lower in *tlr2*-deficient T cells exposed to heat-killed *E. lenta* relative to wild-type controls (**Figure 5C**).

**Figure 5.**
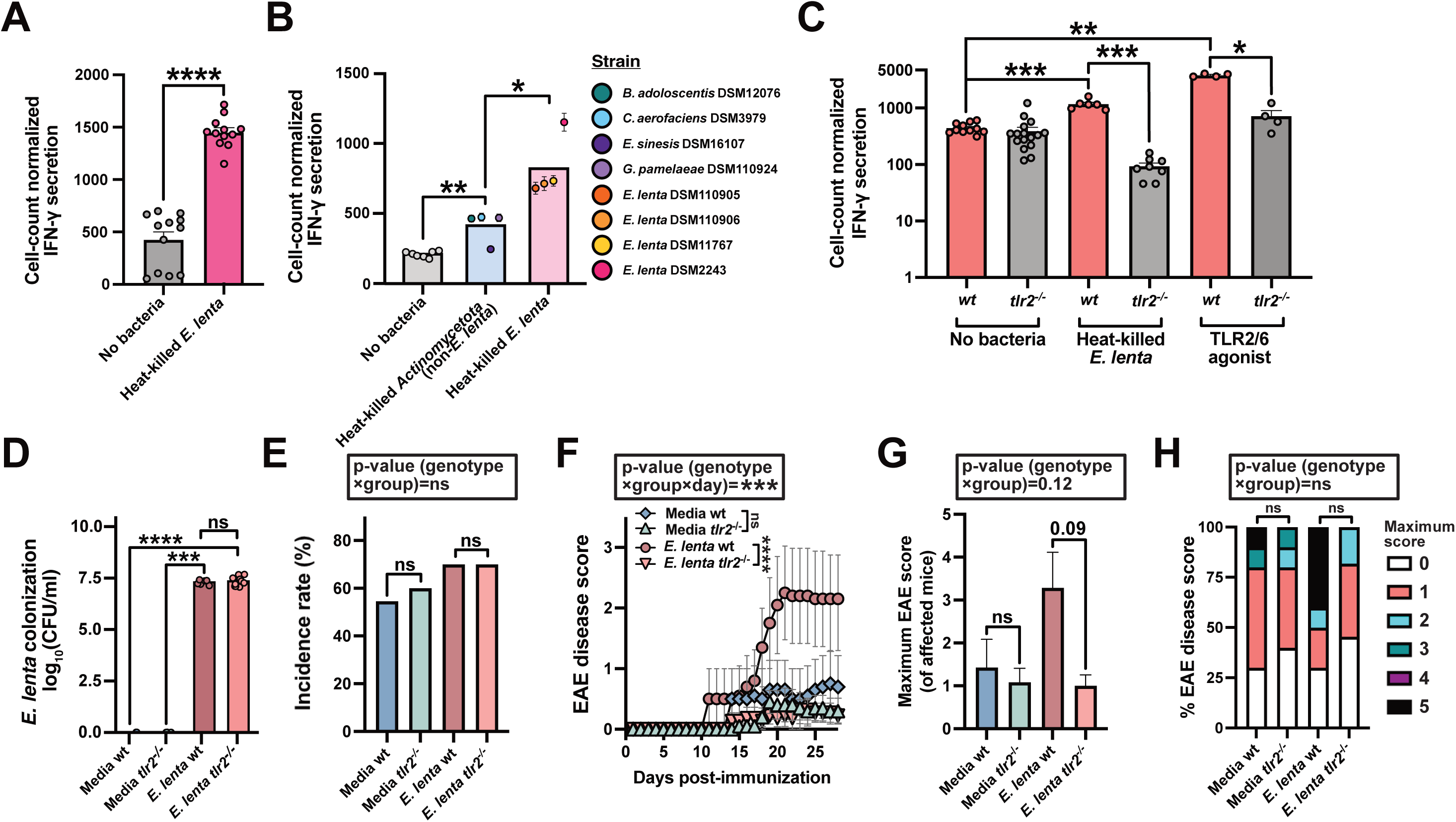
**Induction of IFN-γ and exacerbation of EAE by *E. lenta* requires TLR2.** (A-C) IFN-γ quantification by ELISA from helper T cells cultured with heat-killed bacteria under Th1-skewing conditions. (A) *E. lenta* significantly increases IFN-γ in wt mice (n = 12 replicates/group from 2 experiments). (B) *E. lenta* strains induce stronger responses than other Actinomycetota (n = 7-8 replicates/strain). (C) Induction is significantly reduced in *tlr2^-/-^* T cells (n = 4-16 replicates/group). p values are Wilcoxon rank sum tests (with FDR correction for B, C). (D) Endpoint fecal levels of *E. lenta* as measured by plating on selective media (n=6-11 replicates/group). (E-H) EAE phenotypes in wt and *tlr2^-/-^*mice gavaged with media or *E. lenta* (n = 10-11 mice/group). (E) Incidence, (F) disease score, (G) maximum severity, and (H) peak score proportions. p values are likelihood ratio tests (E), mixed-effects models (F), ANOVA of aligned rank transform [ART] models with Wilcoxon pairwise comparisons (G), or ordinal logistic regression and Fisher’s exact tests (G). Values are mean + SEM (A, C, G), mean (B), percent (E, H), or mean ± SEM (F). *p < 0.05, **p < 0.01, ***p < 0.001, ****p < 0.0001. See also Figures S8-S11.

We tested the hypothesis that *E. lenta* immune activation via host receptor TLR2 mediates its ability to drive neuroinflammation directly using our C57BL/6J EAE model. We repeated oral gavage of *E. lenta* or media control in *tlr2^-/-^* and wt mice, scoring animals over the course of EAE disease (**Figure 5D-H**). *E. lenta* levels were comparable in gavaged mice of *tlr2^-/-^*and wt genotypes (**Figure 5D**) and EAE incidence was comparable between all groups and genotypes (**Figure 5E**). Disease severity significantly differed by genotype for only *E. lenta*-treated mice, with the disease phenotype observed in wt mice completely abrogated in *tlr2^-/-^* animals **(Figure 5F)**. The interaction term between genotype, group, and day was significant, indicating that the CNS inflammatory effect of *E. lenta* is contingent upon host TLR2 signaling **(Figure 5F)**. TLR2 deficient animals also exhibited a downward shift in maximum disease score **(Figure 5G)** and an apparent reduction in severe EAE disease outcomes by score distribution (**Figure 5H**). These results indicate that the ability of *E. lenta* to exacerbate EAE is specific to its immune induction of host receptor TLR2. Taken together, our data support a model in which *E. lenta*, via direct engagement of host TLR2, acts as a critical driver of disease exacerbation, acting as a potent stimulus for (neuro)inflammation once the immune system has been primed by Th1-skewing stimuli such as Complete Freund’s Adjuvant (CFA) and pertussis toxin.

To assess the physiological relevance of our findings further and outside of a disease context, we tested the effect of heat-killed *E. lenta* on T helper signature immune response signature in CONV-R mice, in the context of a complex gut microbiota. As previously demonstrated^24,25^, live *E. lenta* was sufficient to robustly induce Th17 response in ileal lamina propria of conventionally-raised mice (**Figure S11A-D**). Strikingly, heat-killed *E. lenta* also evoked a robust Th17 response (**Figure S11A-D**). Both live and heat-killed *E. lenta* further induced robust expression of Th1 master transcription factor TBET (**Figure S11E-F**). We did not observe any changes in expression of regulatory T cell marker FoxP3 (**Figure S11G-H**). In sum, our results suggest that innate recognition of *E. lenta* cell components by TLR2 promotes Th1 and Th17 differentiation, providing a cellular mechanism through which *E. lenta* predisposes to immune phenotypes such as EAE.

## DISCUSSION

Taken together, our results suggest that the human gut microbiota is an underappreciated contributor to sex disparities in autoimmune disease risk. We revealed a robust association between human gut microbiota composition and sex in high-quality metagenomic datasets from dozens of independent studies^19^. Further, by re-analyzing data from the largest human study of the microbiome in multiple sclerosis^21^, we discovered that species most enriched in women, but not men, positively associate with MS status and severity. We went on to provide causal evidence that one of the top female-enriched species, *E. lenta*, drives neuroinflammation in mice, exacerbating disease phenotypes and T cell infiltration into the brain. These effects were independent of a previously-identified gene responsible for Th17 activation^25^ and consistent with the ability of *E. lenta* to robustly induce the Th1 signature cytokine IFN-γ in healthy mice and cell culture assays. IFN-γ production relied on sensing of *E. lenta* cell components via TLR2.

We opted to focus on *E. lenta* given the extensive tools for studying this bacterial species as a model Actinomycetota^23,24,29^ and a rich prior literature associating *E. lenta* with MS risk and severity^15,30–32^. However, many other sex-associated gut bacteria identified in this study have potential impacts on MS. This includes *E. tayi*^14,21^, the most significantly enriched species in female subjects, and *Akkermansia muciniphila*^7,8,16,30,31,33^, which can exacerbate EAE: both species are associated with human MS. Female-enriched *Clostridium leptum*^34,35^ is also positively associated with MS disease and its severity. Notably, while causal evidence in mice remains outstanding, prior literature on the identified male-enriched bacteria support a potential protective role in MS: top male-enriched bacterial species by abundance, *Segatella* [formerly *Prevotella*] *copri*^30,32,36^, as well as *Bifidobacterium adolescentis*^14^ and *Slackia isoflavoniconvertens*^33^ are all depleted in PwMS. Thus, female bias of MS may be driven by increased exposure to pathogenic bacterial species combined with decreased levels of beneficial bacteria.

Our data strengthens recent research indicating that despite its prevalence in the gut microbiota of healthy humans, *E. lenta* can be pathogenic across multiple disease contexts, representing a potential “pathobiont”. Traditionally, *E. lenta* was only implicated in bacteremia^37^; however, recent studies have provided causal evidence that *E. lenta* can exacerbate inflammatory bowel disease^25^ and rheumatoid arthritis^25,38^. Herein, in both mice otherwise lacking gut microbiota and in conventionally reared mice, we show that *E. lenta* aggravates the EAE model and is sufficient to induce both Th1 and Th17 immune response in otherwise healthy mice. *E. lenta* provides a promising model to develop further mechanistic insight into the role of the gut microbiota in autoimmunity due to its ease of growth^23^ and recently-developed genetic tools^24,39^, in contrast to the widely-studied but microbiologically intractable mouse-derived segmented filamentous bacteria (SFB)^40^. Furthermore, the high prevalence and reproducible disease associations of *E. lenta* in humans supports the translational relevance of this line of inquiry.

Surprisingly, we found that the impact of *E. lenta* on the EAE model was independent of the *cgr* operon, which we previously showed was sufficient to mediate colonic, but not ileal, Th17 response and to exacerbate colitis in mice^24,25^. In EAE, both wt and Δ*cgr E. lenta* strains induced colonic and brain Th responses, driving robust disease phenotypes. Interestingly, the impact of classic EAE inducer SFB on Th17 cells is restricted to the ileum, which is sufficient to exacerbate EAE but, without local colonic induction or strong Th1 engagement, results in mixed or even protective effects in colitis^6,41–44^. Our data supports a broader impact of *E. lenta* on the intestinal immune system, leading to a robust ileal and colonic Th1 signature, consistent with the well-documented contributions of both Th1 and Th17 immune responses to EAE phenotypes^6,9–12^ and human MS^45,46^. The impact of *E. lenta* on host immunity may also extend beyond T cells, as recent work suggests altered IgA recognition of *E. lenta* by B cells in _PwMS_^4^7,4^8^.

Our combined observations *in vitro* and *in vivo* indicate that *E. lenta* potently activates TLR2, leading to strain-variable increases in IFN-γ production by T helper cells in the presence of Th1 cytokines. TLR2 signaling within helper T cells has been shown to directly modulate EAE severity, independent of antigen-presenting cells, by shaping pro-inflammatory effector programs driving disease ^26,49^. Consistent with this framework, during EAE we observed a robust induction of IFN-γ by *E. lenta* in both gut lamina propria and CNS, alongside a more limited Th17 response. These data support a model in which innate sensing of *E. lenta* through TLR2 biases intestinal T cell differentiation toward pro-inflammatory Th1-associated programs that can exacerbate neuroinflammation upon disease induction. Interestingly, a recent report identified a pro-inflammatory lipid in *E. lenta* DSM2243 that signals via TLR2^50^, though given this lipid’s requirement for electrophilic activation, a different, distinct ligand is likely involved in *E. lenta*’s ability to drive Th1 and EAE.

Ultimately, our work emphasizes the importance of considering sex as a biological variable, not just as a confounder but as a potential driver of microbiome-dependent phenotypes. While host sex is routinely captured in human microbiome association studies, it has rarely been the focus of any downstream experimental efforts. This is due in part to lack of studies like this that provide a proof-of-concept for studying sex-associated bacteria coupled to a clear clinical motivation for these studies–the alarming and increasing prevalence of MS in women relative to men. A major open question is the environmental and/or genetic factors that reinforce sex differences in the gut microbiota, which would be invaluable in designing better preclinical models to mimic the differences in gut microbial community structure between male and female human subjects. Continued progress in this area promises to address long-standing disparities in biomedical science and medical practice through a more comprehensive microbiome-informed view of sex disparities in disease risk, progression, and treatment outcomes.

### Limitations of this study

While we identify a robust sex-signature in the human gut microbiota, relative contributions of sex hormones versus chromosomal factors as drivers remain undetermined. This presents an interesting line of inquiry, especially considering recent research demonstrating *E. lenta* metabolizes sex hormones and other steroids, with immune relevance^39,51^. Additionally, while our meta-analysis revealed striking associations between MS disease and numerous sex-linked microbes, many of which have been indirectly linked to neuroinflammation, *E. lenta* represents the only female-enriched human pathobiont shown to independently drive disease in mice. Indeed, while strains of *Akkermansia* have been implicated in EAE, as previously discussed, prior evidence was limited to mouse-derived isolates. Finally, while we have determined that *E. lenta* acts to induce both EAE and robust, strain-variable IFN-γ response via host receptor TLR2, further work is warranted to identify the specific bacterial ligand(s) responsible for this effect.

## RESOURCE AVAILABILITY

The resource availability section is required for all research articles. This section might also be required for applicable reviews and perspectives. This section must contain the following required subsections under the resource availability heading: “lead contact,” “materials availability,” and “data and code availability.”

For complete information on requirements, including formatting instructions, refer to the Info for Authors page specific to the journal you are publishing with.

## Lead contact

Requests for further information and resources should be directed to and will be fulfilled by the lead contact, Peter J. Turnbaugh.

## Materials availability

This study did not generate new unique reagents.

## Data and code availability

● This paper analyzes existing, publicly available data.
● The human metagenomic data were accessed via the curatedMetagenomicData R package (v3.16). This database was analyzed using an openly available custom script.
● The iMSMS clinical dataset is publicly available at DOI: 10.1016/j.cell.2022.08.021.
● RNA sequencing data for mouse CD4^+^ T cells were obtained from (Alexander et al., 2021, DOI: 10.1016/j.chom.2021.11.001) and are available upon request.
● The interactive meta-analysis volcano plots are provided as Supplementary Data 1 and Supplementary Data 2.
● This paper does not report original code.
● Any additional information required to reanalyze the data reported in this paper is available from the lead contact upon request.

## ACKNOWLEDGEMENTS

We thank the members of the Turnbaugh lab for their helpful feedback throughout this project and we thank Dr. Matthew Spitzer’s lab for sharing *in vitro* assay troubleshooting tips and insight, with specific appreciation for Camille Valente and Janice Arakawa-Hoyt. We also thank the UCSF Gnotobiotics Core (gnotobiotic mouse experiments); the Scharschmidt lab (*tlr2^-/-^* mice); the Balskus lab (Δ*cgr E. lenta* and *wild-type* control strain); the UCSF Flow Cytometry Core (P30DK063720); the UCSF Institute for Human Genetics core (RNA-seq); and the UCSF Media Production and Antibody Core (media). Funding was provided from the National Institutes of Health (R01CA255116, R01DK114034, and R01HL122593 to P.J.T.) as well as the Swiss National Science Foundation (SNF Eccellenza Professorship: PCEFP3_194609; SNF Starting Grant: TMSGI3_211318), the National MS Society (FG-1708-28871), the Propatient Foundation, and the Gottfried and Julia Bangerter Rhyner Foundation (all to A.-K.P). Additionally, fellowship support to RRR was received from the National Institutes of Health UCSF T32 Endocrine Training Grant (Grant No. 5T32DK007418), and the National Science Foundation Graduate Research Fellowship Program (Grant No. 2038436). Any opinions, findings, and conclusions or recommendations expressed in this material are those of the author(s) and do not necessarily reflect the views of the National Science Foundation. S.E.B. holds the Heidrich Friends and Family endowed Chair at UCSF Neurology. P.J.T is a Biohub Investigator. Finally, we would like to acknowledge the Ramaytush Ohlone people, who are the traditional custodians of this land. We pay our respects to the Ramaytush Ohlone elders, past, present, and future, who call this place, the land that UCSF sits upon, their home.

## AUTHOR CONTRIBUTIONS

Conceptualization, R.R.R. and P.J.T.; methodology, R.R.R., P.J.T., M.A., C.N., K.T., V.U., L.R., L.S., and A.K.P.; investigation, R.R.R., M.A., E.O., L.R., K.T., C.O., L.S., and T.H.; validation, R.R.R., E.O., and L.R.; visualization, R.R.R., P.J.T., and E.O.; writing—original draft, R.R.R. and P.J.T.; writing—review & editing, all authors; funding acquisition, R.R.R. and P.J.T.; resources, A.K.P.; supervision, P.J.T., A.K.P., and S.E.B.; project administration, P.J.T.

## DECLARATION OF INTERESTS

P.J.T. is on the scientific advisory boards of Pendulum and SNIPRbiome. All other authors declare no competing interests.

## DECLARATION OF GENERATIVE AI AND AI-ASSISTED TECHNOLOGIES

AI was used for proofreading for frank typos. Generative AI and AI-Assisted technologies were not used for any other purposes.

## SUPPLEMENTAL INFORMATION

Document S1. Figures S1-S12

Document S2. Tables S1-S10

Supplementary Data 1. Interactive volcano plot for sex-associated species abundance model, related to **Figure 1** and Figures S1 and S2

Supplementary Data 2. Interactive volcano plot for sex-associated species prevalence model, related to **Figure 1** and Figure S1

## SUPPLEMENTARY FIGURE LEGENDS

**Figure S1.**
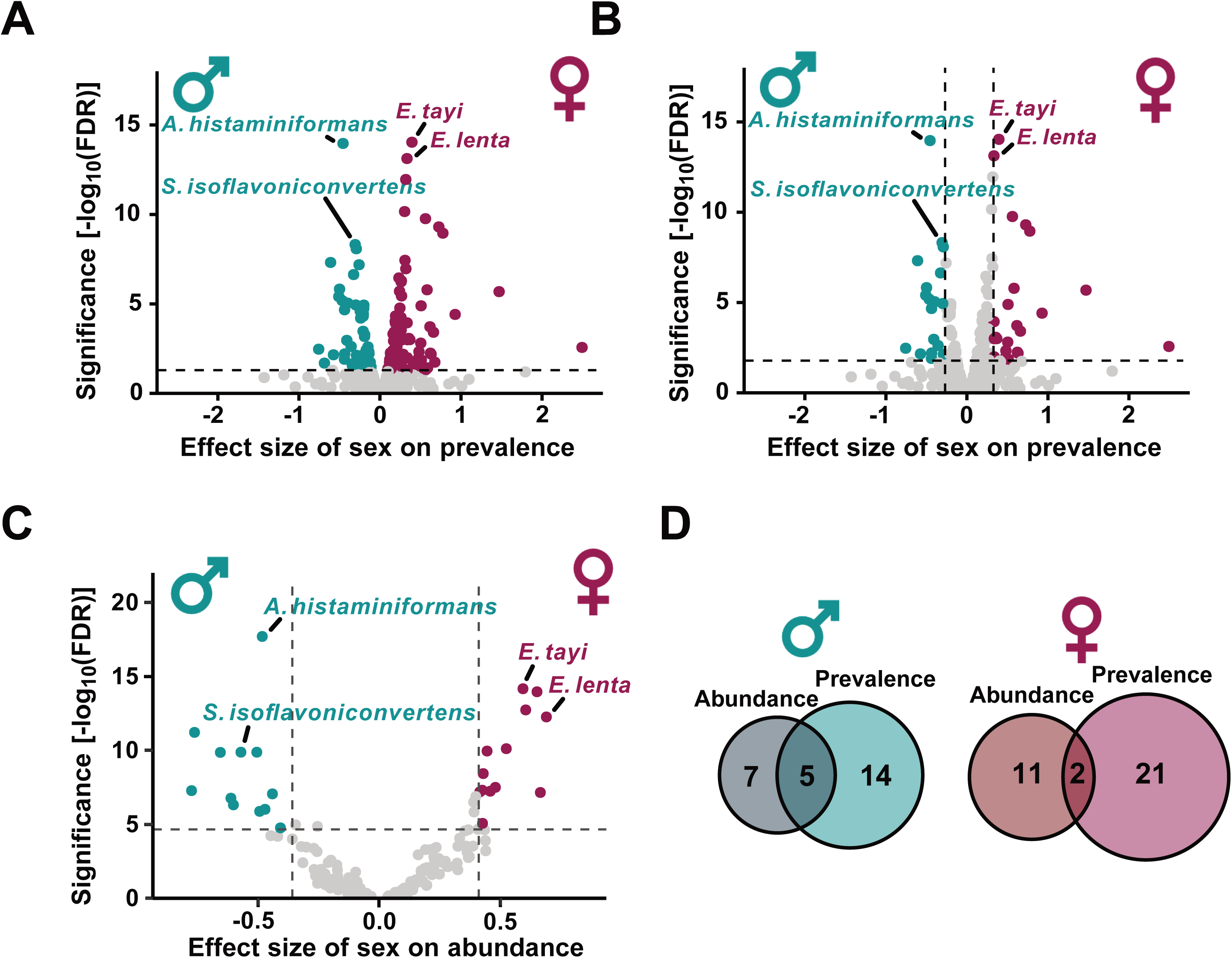
Meta**-analysis reveals sex-associated gut bacterial species, related to Figure 1**. (A) Differentially prevalent microbial species by sex in the curatedMetagenomicData database (dashed line indicates FDR < 0.05) determined via FDR-adjusted logistic models (glm(species prevalence ∼ sex + age + BMI + continent + health + median read length + sequencing depth + DNA extraction kit, family=binomial)). (B, C) Top hits using a stricter cutoff (dashed lines indicate lowest quintile FDR and highest quintile effect size) for (B) prevalence and (C) abundance. (B) Logistic model with with sex, age, BMI, continent, health, and sequencing metrics as fixed effects (species prevalence ∼ sex + age + BMI + continent + health + median read length + sequencing depth + DNA extraction kit, family=binomial). (C) MaAsLin2 model with sex, age, BMI, continent, health, and sequencing metrics as fixed effects and study as a random effect (CLR-normalized relative abundance ∼ sex + age category + BMI category + continent + health category + median read length category + sequencing depth category + DNA extraction kit category + (1|study)). (D) Venn diagrams display the top sex-associated bacterial species common to both (B) prevalence and (C) abundance models. (A-C) Each dot represents one species, colored by sex of enrichment.(A) Differentially prevalent microbial species by sex in the curatedMetagenomicData database (dashed line indicates FDR < 0.05). (B, C) Top hits using a stricter cutoff (dashed lines indicate lowest quintile FDR and highest quintile effect size) for (B) prevalence and (C) abundance. (D) Venn diagrams displayed the top sex-associated bacterial species common to both (B) prevalence and (C) abundance models. (A-C) Each dot represents one species, colored by sex of enrichment. Statistics: (A, B) FDR-adjusted logistic model (glm(species prevalence ∼ sex + age + BMI + continent + health + median read length + sequencing depth + DNA extraction kit, family=binomial), STAR Methods), (C) FDR-adjusted MaAsLin2 model (CLR-normalized abundance ∼ sex + age category + BMI category + continent + health category + median read length category + sequencing depth category + DNA extraction kit category + (1|study), STAR Methods).

**Figure S2.**
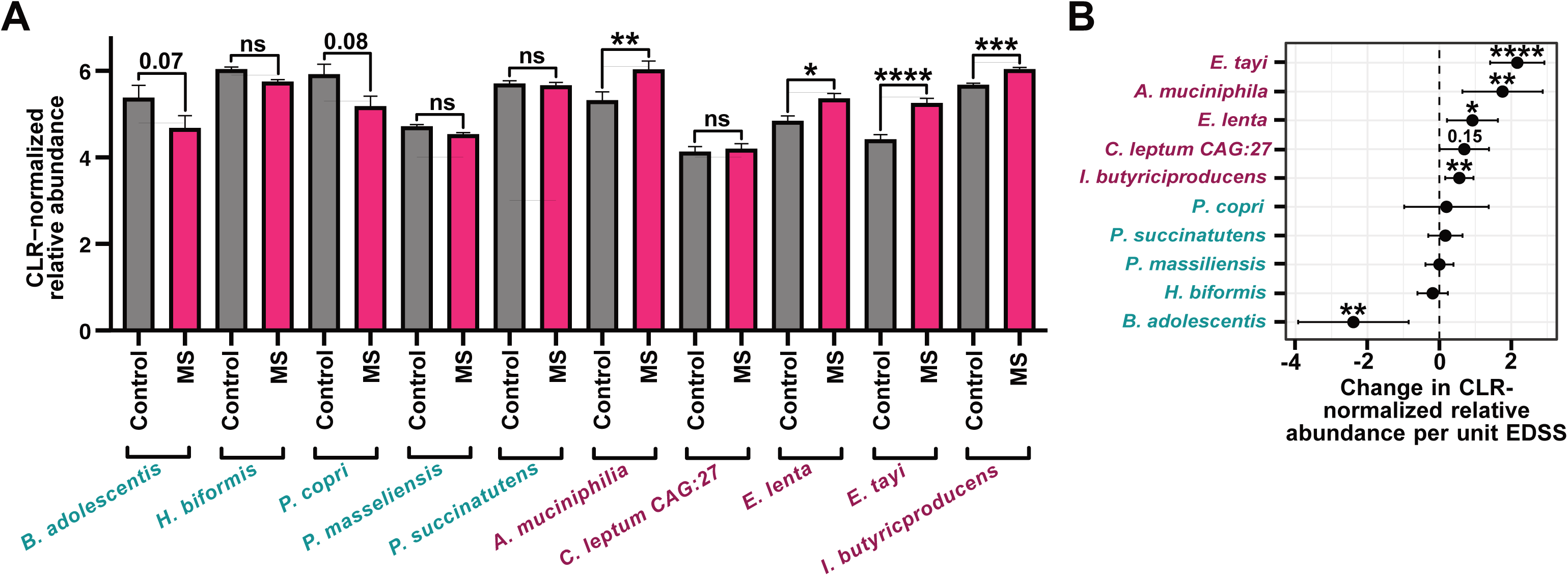
Sex-linked bacterial species association with MS is detectable using an alternative selection criterion, related to. **Figure 1**. The top 10 differentially abundant species by effect size (Figure 1A, 5 female, 5 male) were analyzed in the iMSMS dataset. (A) The CLR-normalized relative abundance of 4/5 female-enriched species detected were significantly enriched in stool samples from people living with MS (PwMS) relative to healthy controls. (B) All 5 female-enriched species were associated with disease severity (Expanded Disability Status Scale [EDSS] score). p values for (A, B) were determined via sex- and treatment-adjusted linear models: CLR-normalized relative abundance ∼ disease status + sex + treatment status; CLR-normalized relative abundance ∼ EDSS score + sex + treatment status. *p < 0.05, **p < 0.01, ***p < 0.001, and ****p < 0.0001. Full species names: *Akkermansia muciniphila, Bifidobacterium adolescentis, Clostridium leptum CAG:27, Eggerthella lenta, Eisenbergiella [Dialister] tayi, Holdemanella biformis, Intestinimonas butyriciproducens, Phocea [Phocaeicola] massiliensis, Prevotella [Segatella] copri,* and *Phascolarctobacterium succinatutens*.

**Figure S3.**
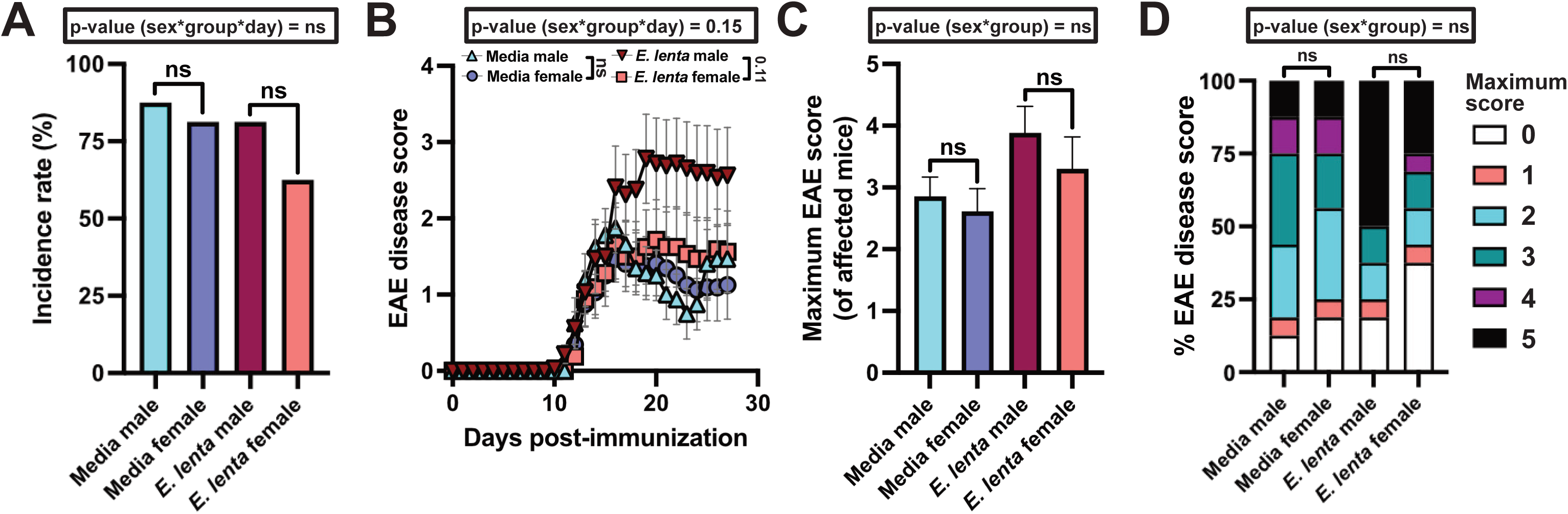
**Sex does not significantly alter disease severity in media or *E. lenta*-treated mice.** Mice were gavaged with media or *E. lenta* DSM2243 starting one week prior to induction of disease (Figure 2A). (A-D) EAE phenotypes were tracked for: (A) EAE incidence, determined by likelihood ratio tests of a global generalized linear model with binomial error and logit link (glm(survival ∼ group * sex, family = binomial) and subset models; (B) EAE disease score, analyzed via repeated measures mixed-effects model with mouse as a random effect (response variable [EAE disease score] ∼ sex * group * day + (1|mouse)); (C) maximum disease severity, assessed by ANOVA of Aligned Rank Transform (ART) model (score ∼ group * sex) with Wilcoxon rank sum test for pairwise comparisons; and (D) peak score proportions, evaluated by ordinal logistic regression and Fisher’s exact test. Values are (B) mean ± SEM, (C) mean + SEM, and (A, D) percentage.

**Figure S4.**
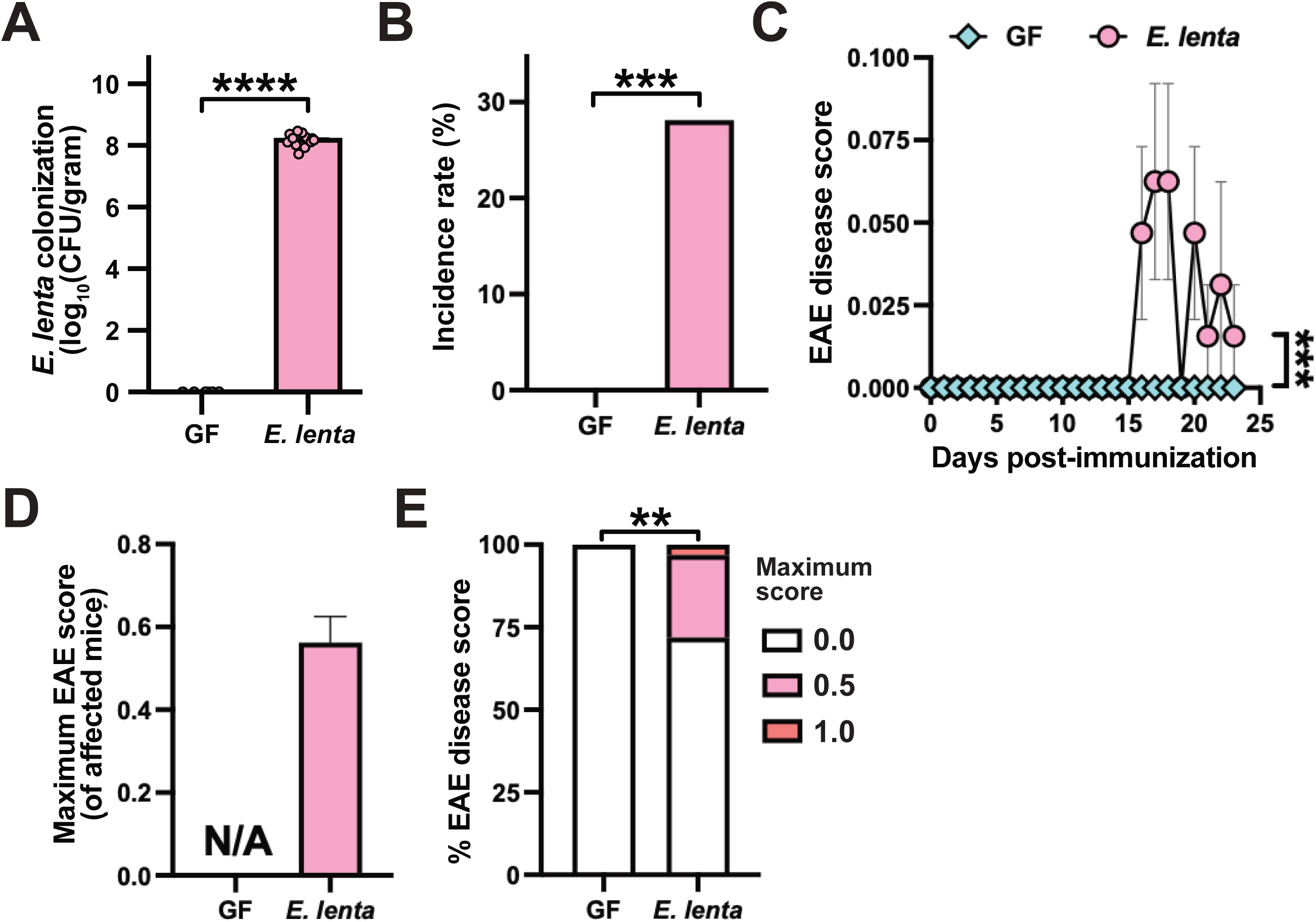
***E. lenta* is sufficient to induce mild EAE disease phenotypes in GF mice.** Male (n = 12 mice/group) and female (n = 24 mice/group) germ-free (GF) mice were colonized with *E. lenta* DSM2243 or no bacteria starting 2-4 weeks prior to disease induction. (A) Viable *E. lenta* levels (log_10_ CFU/gram) measured via plating; p values are Wilcoxon rank sum tests. (B-E) EAE phenotypes were tracked for: (B) incidence, using likelihood ratio tests of a global generalized linear model (glm(incidence ∼ group, family = binomial)); (C) disease score, using repeated measures mixed-effects models (response variable ∼ group * day + (1|mouse)); (D) maximum disease severity, using Wilcoxon rank sum tests; and (E) peak score proportions, using Fisher’s exact tests. Values are presented as (A, D) mean + SEM, (B, E) percentage, (C, D) mean ± SEM. **p < 0.01, ***p < 0.001, and ****p < 0.0001.

**Figure S5.**
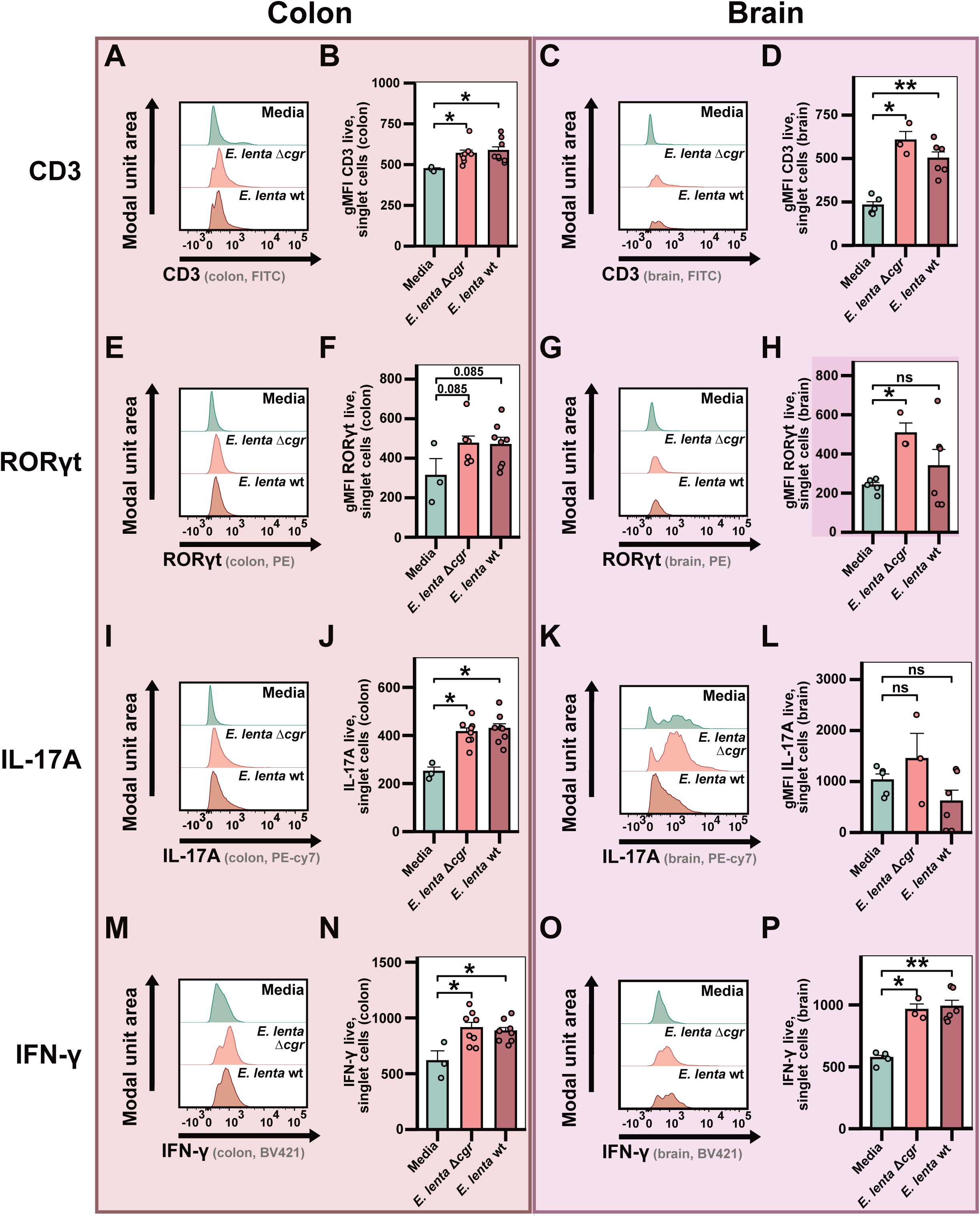
***E. lenta* induces local immune response in colonic lamina propria and remote response in the brain during EAE, independent of the *cgr* operon.** Peak disease (day 15 post-immunization) flow cytometry data from the colonic lamina propria and brains of adult, male mice gavaged with media, *Δcgr E. lenta* DSM2243, or wt *E. lenta* DSM2243 (n = 3-8/group; see Tables S7, S8). (A, C, E, G, I, K, M, O) Representative histograms. (B, D, F, H, J, L, N, P) Quantification of gMFI among live, singlet cells for: CD3, RORγt, IL-17A, and IFN-γ. p values are Wilcoxon rank sum tests. *p < 0.05 and **p < 0.01.

**Figure S6.**
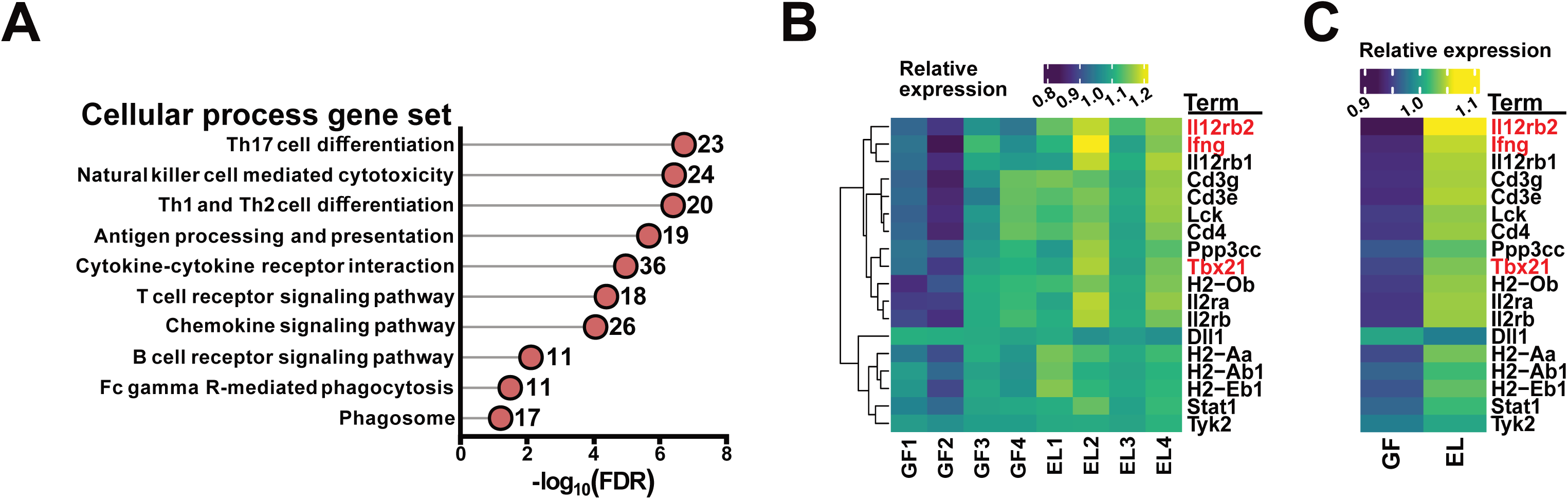
***E. lenta* monocolonization induces robust transcriptional response in ileal helper T cells.** Pathway enrichment analysis of published RNA-seq data from ileal helper T cells (n = 4 mice/group). DEGs were identified using DESeq2 (|Fold change| > 1.5; Wald test p < 0.05). (A) Top ten hits for cellular process terms, analyzed via Fisher’s exact test (FDR shown). (B, C) Relative expression (VST value) of DEGs from the Th1 and Th2 cell differentiation pathway displayed by (B) sample and (C) group average; core Th1 genes are highlighted in red. See also Tables S9 and S10.

**Figure S7.**
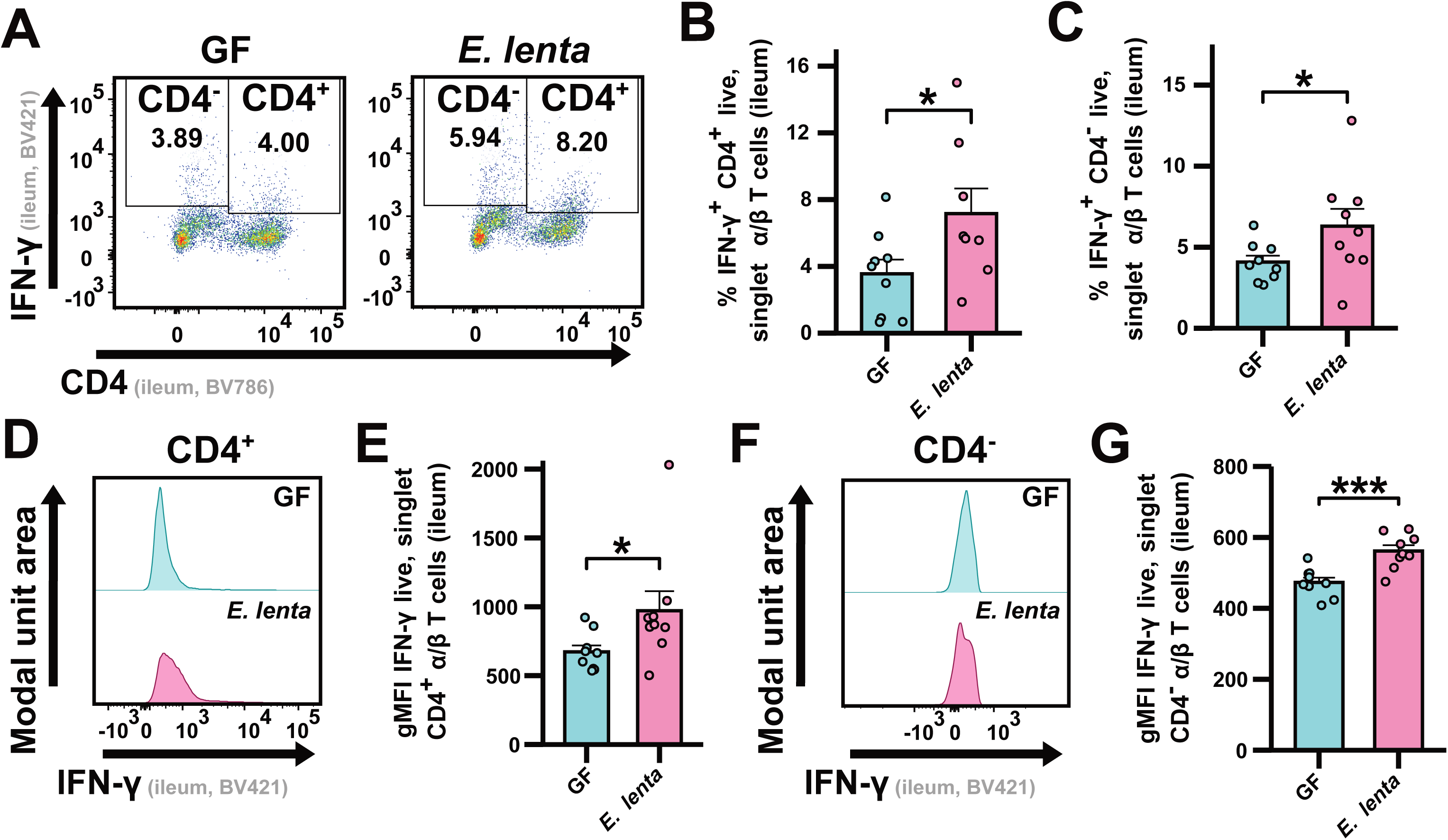
***E. lenta* drives expansion of IFN-γ^+^ CD4^+^ and CD4^-^ cells among ileal T cells in homeostasis, related to Figure 4**. (A-D) Flow cytometry analysis of the ileal lamina propria of E. lenta monocolonized and GF mice (n = 8-9 mice/group). (A) Representative flow plots. (B-C) gMFI IFN-γ of live, singlet Tcr-beta^+^ (α/β) T cells positive (B) or negative (C) for CD4. (D, F) Representative histograms. (E, G) gMFI IFN-γ of live, singlet Tcr-beta^+^ T cells positive (E) or negative (G) for CD4. p values are Wilcoxon rank sum tests. *p < 0.05 and ***p < 0.001.

**Figure S8.**
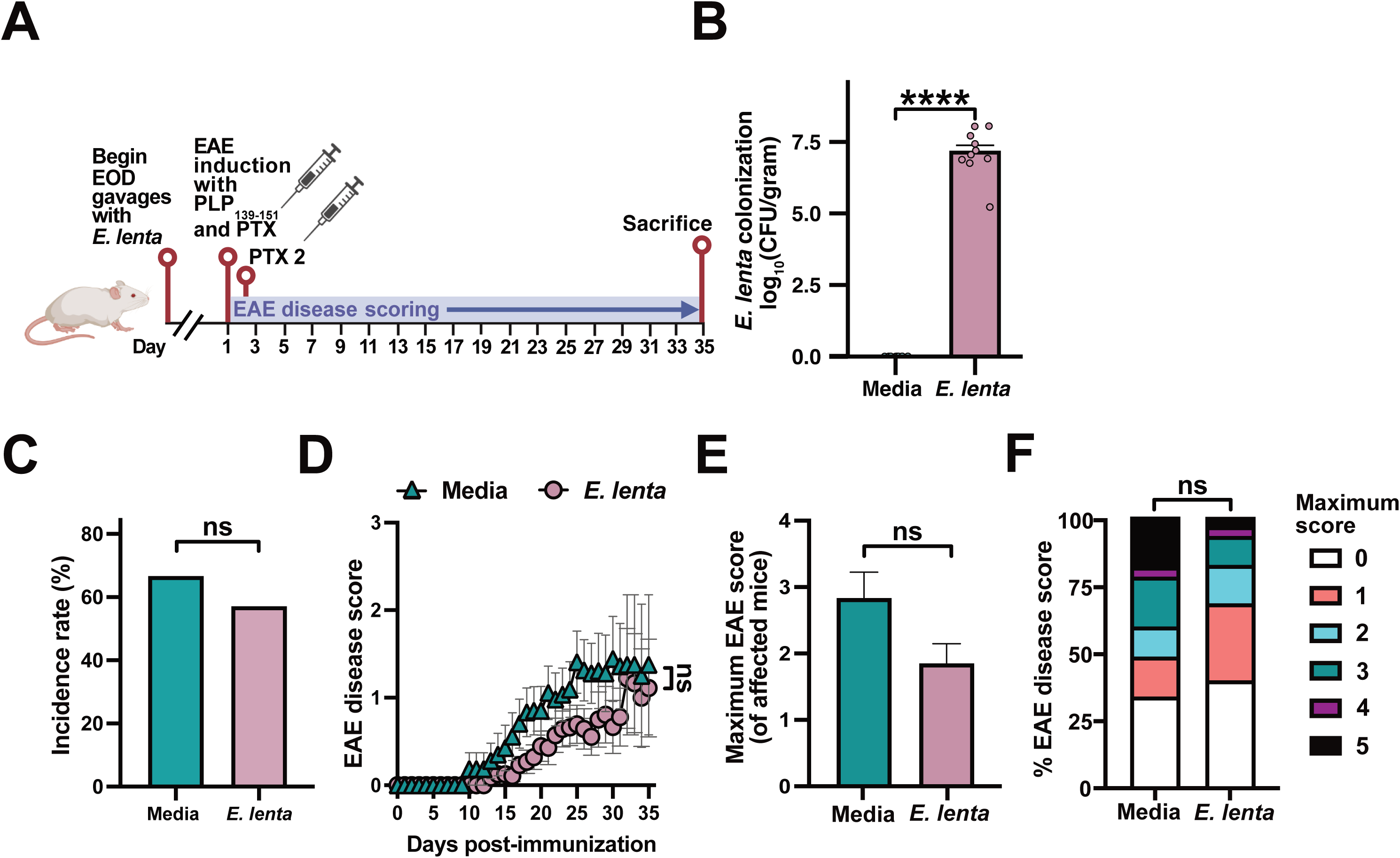
*E. lenta* does not exacerbate an alternative model of EAE, related to Figure 5. (A) SJL/J mice were gavaged with media or *E. lenta* DSM2243 starting two weeks prior to disease induction. (B) *E. lenta* levels (log_10_ CFU/g) measured via qPCR; p values are Wilcoxon rank sum tests. (C-F) EAE phenotypes were tracked for: (C) incidence and (D) disease score, using repeated measures mixed-effects models (response variable ∼ group * day); (E) maximum disease severity, using Wilcoxon rank sum tests; and (F) peak score proportions, using Fisher’s exact test. Values are (B, E) mean + SEM, (C, F) percentage, or (D) mean ± SEM. ****p < 0.0001.

**Figure S9.**
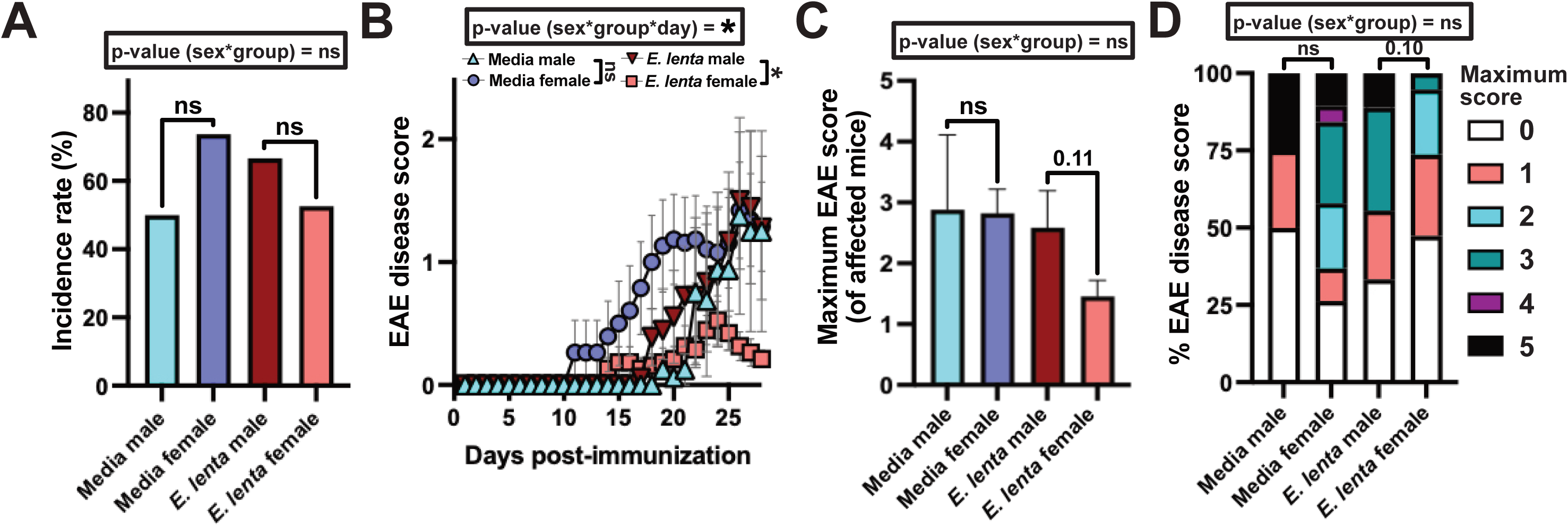
Neuroinflammation is more modest in female mice colonized with *E. lenta* in an alternative model of EAE, related to Figure 5. Mice were gavaged with media or *E. lenta* starting one week prior to disease induction (Figure S8A). (A-D) EAE phenotypes were tracked for: (A) incidence, using likelihood ratio tests of a global GLM; (B) disease score, using repeated measures mixed-effects models (response variable ∼ sex * group * day + (1|mouse)); (C) maximum disease severity, using ANOVA of ART models with Wilcoxon pairwise comparisons; and (D) peak score proportions, using ordinal logistic regression and Fisher’s exact tests. *p < 0.05. Values are (A, D) percentage, (B) mean ± SEM, or (C) mean + SEM.

**Figure S10.**
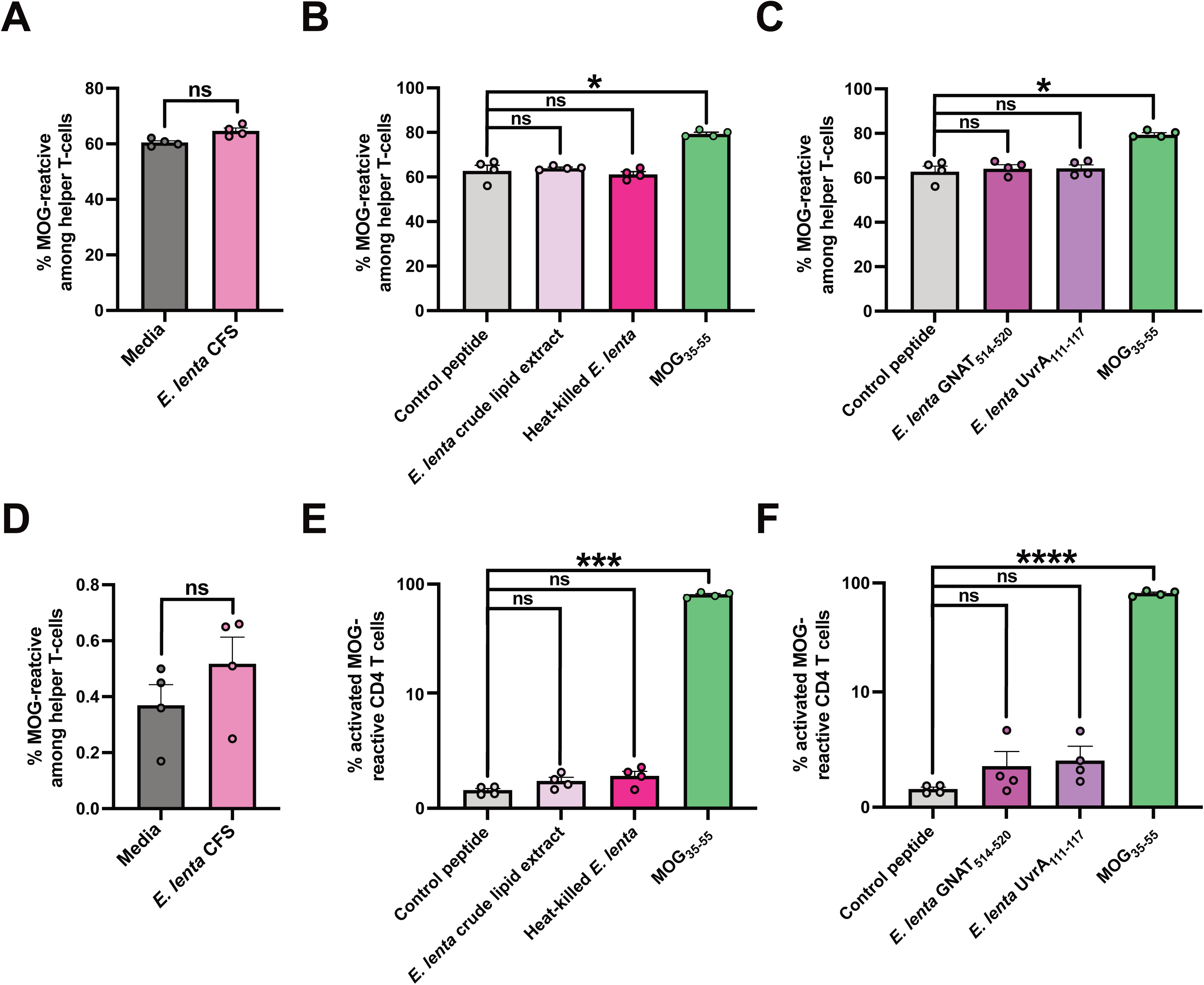
***E. lenta* does not evoke an anti-MOG response, related to Figure 5**. (A-F) Helper T cells and dendritic cells were isolated from 2D2 mice and co-cultured with specified antigens. (A, D) *E. lenta* cell-free supernatant (CFS) (A) and spent media (D) did not expand the MOG-reactive fraction; p values are paired Wilcoxon rank sum tests. (B, C, E, F) Percent activation of MOG-reactive helper T cells was assessed for *E. lenta* cells or peptides compared to MOG_35-55_; p values are FDR-corrected paired Welch’s T-tests. Values are mean + SEM. ns = nonsignificant, *p < 0.05, ***p < 0.01, ****p < 0.001.

**Figure S11.**
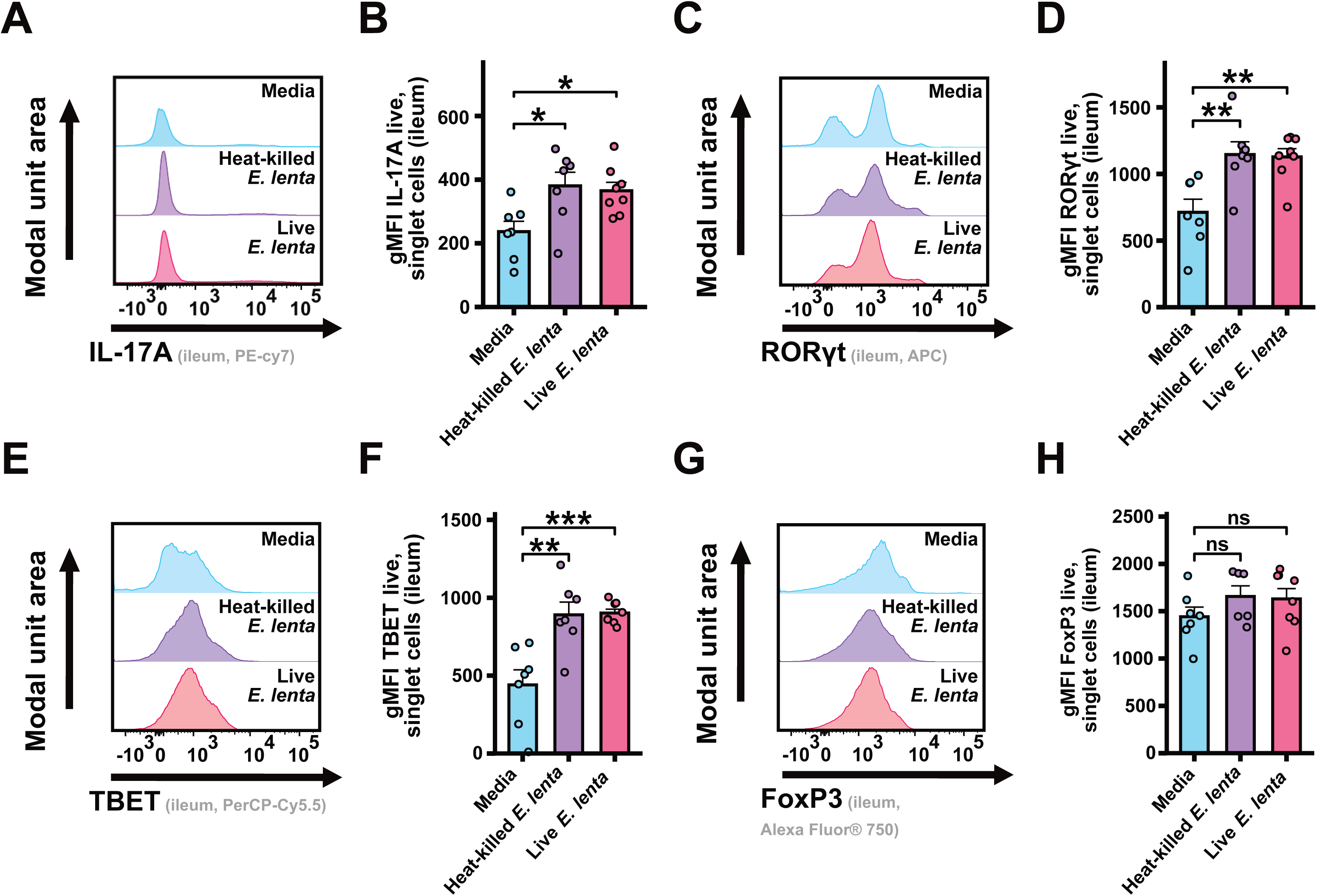
***E. lenta*, independent of viability, induces core immune markers in the ileal lamina propria of healthy CONV-R mice, related to Figure 5**. Mice were gavaged with media, heat-killed *E. lenta*, or live *E. lenta* (n = 7-8/group). (A, C, E, G) Representative histograms. (B, D, F, H) Quantification of gMFI among live, singlet-cells for: IL-17A, RORγt, TBET and FoxP3. p values are Wilcoxon rank sum tests. *p < 0.05, **p < 0.01, ***p < 0.001. Values are mean + SEM.

**Figure S12.**
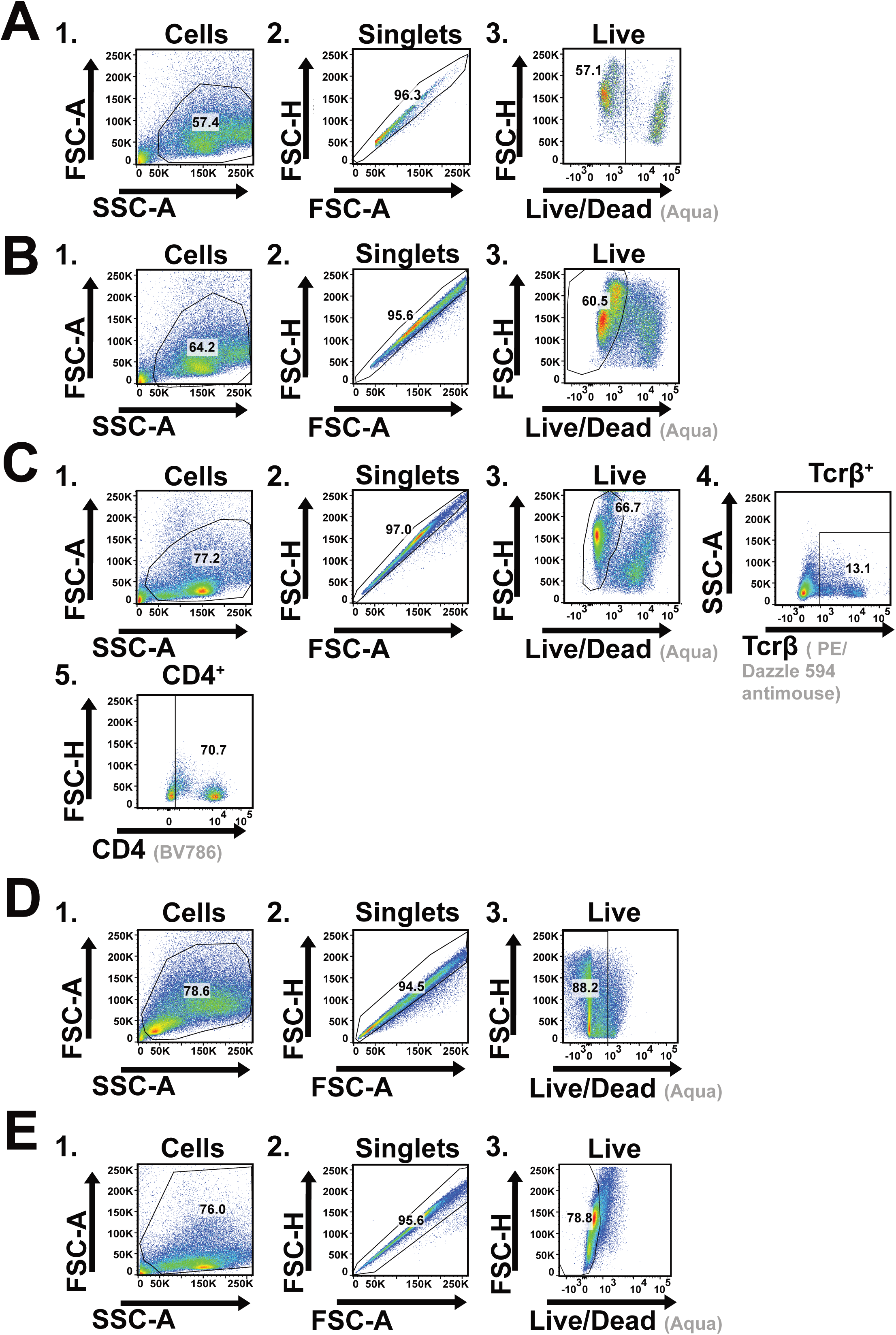
**Flow cytometry gating strategies, related to STAR Methods.** All flow gates are sequential; numbers indicate application order. (A, B) Brain and colonic lamina propria data from mice colonized with wt or *Δcgr E. lenta* (see Figure 3 and Figure S5). (C, D) Ileal and colonic lamina propria data from GF or *E. lenta* monocolonized mice (see Figure 4 and Figure S7). (E) Ileal lamina propria data from mice treated with live or heat-killed *E. lenta* (see Figure S11).

## STAR★METHODS

### KEY RESOURCES TABLE

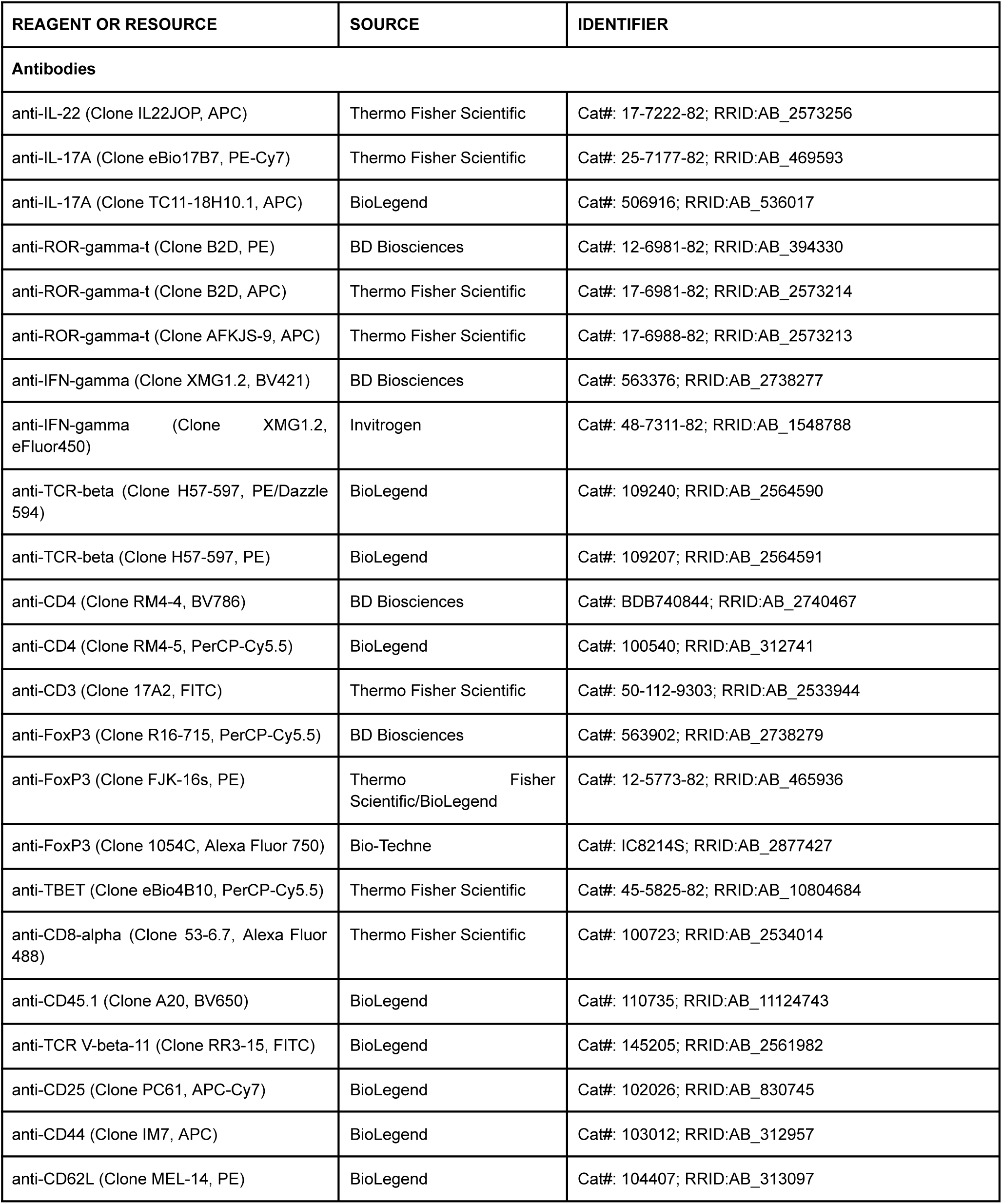
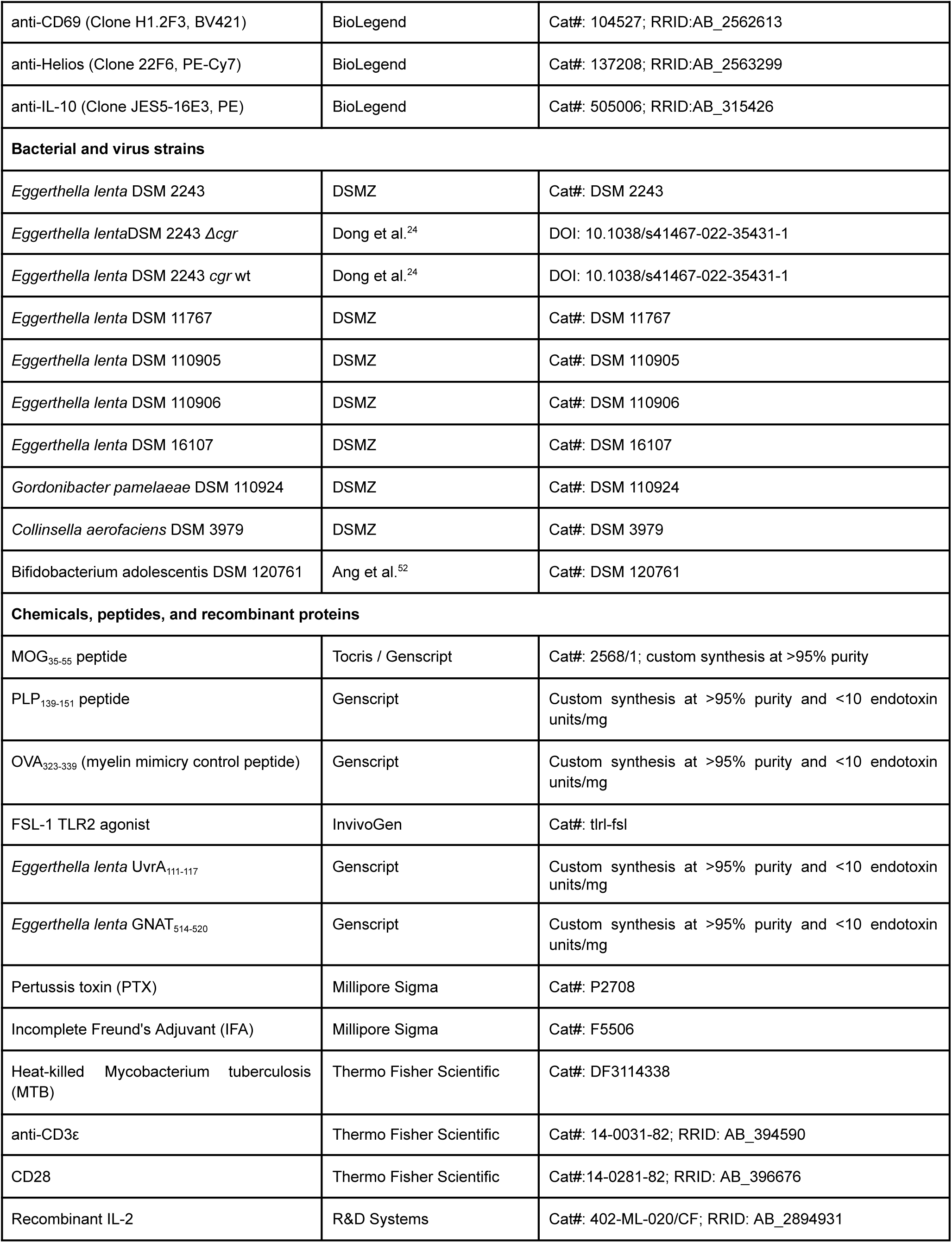
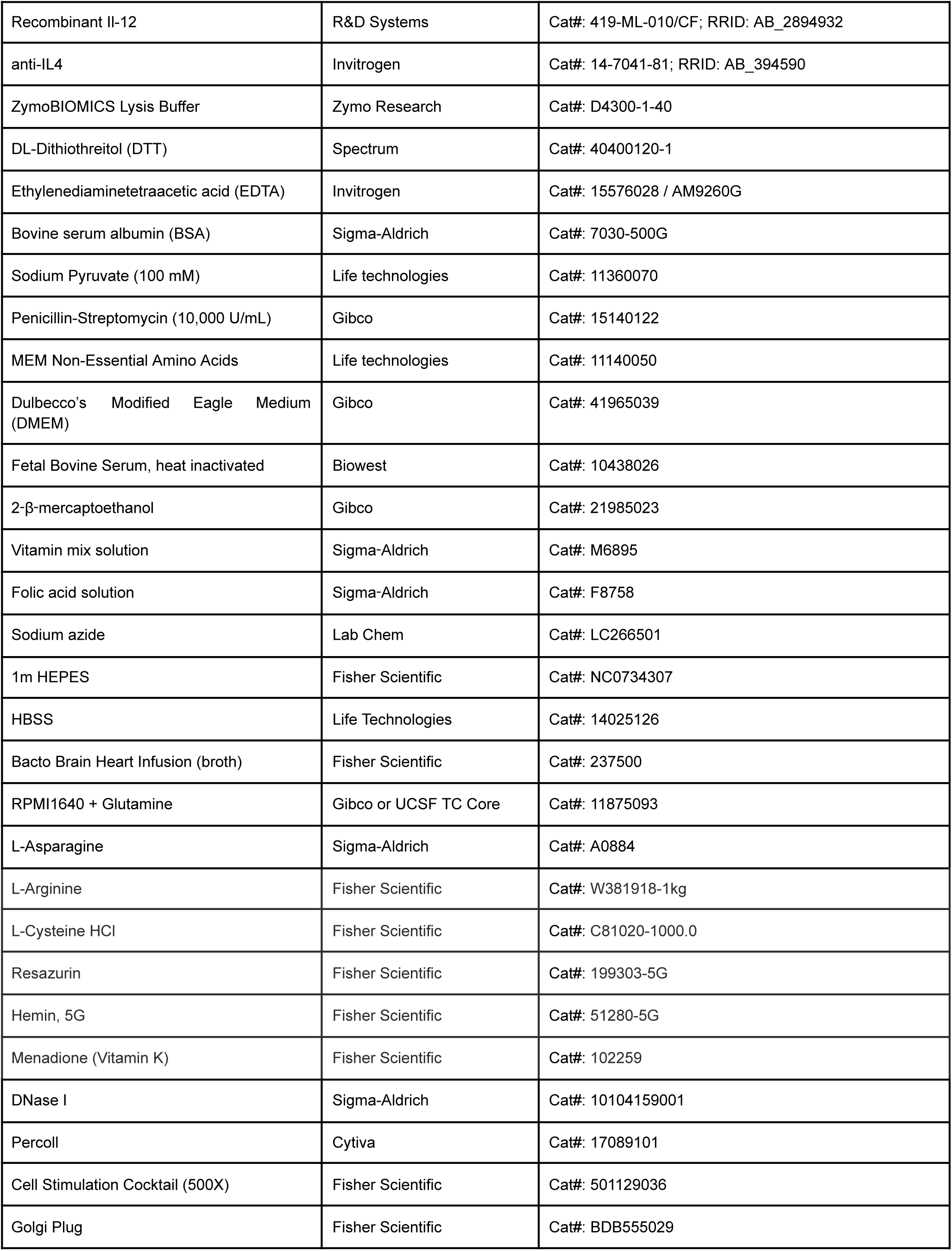
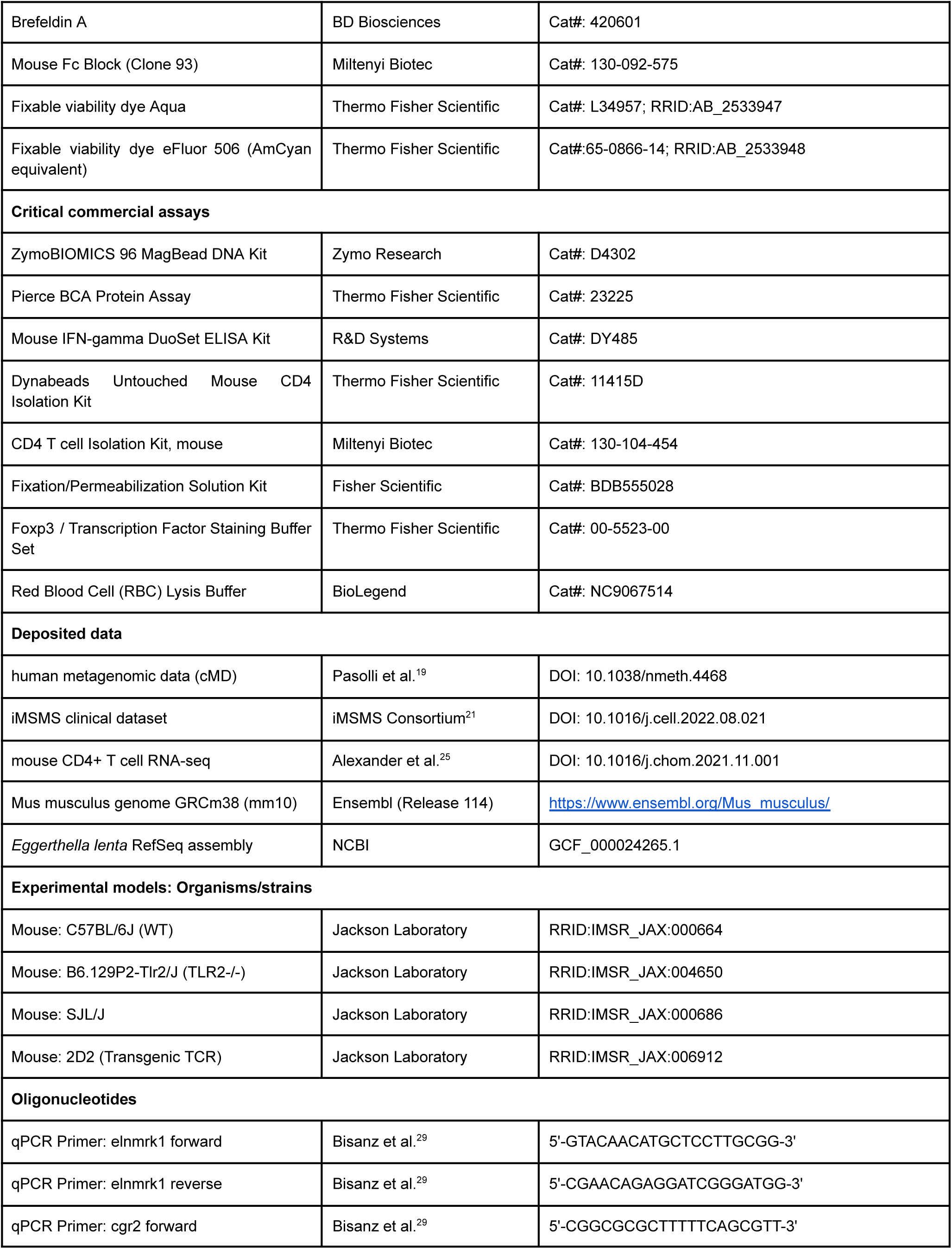
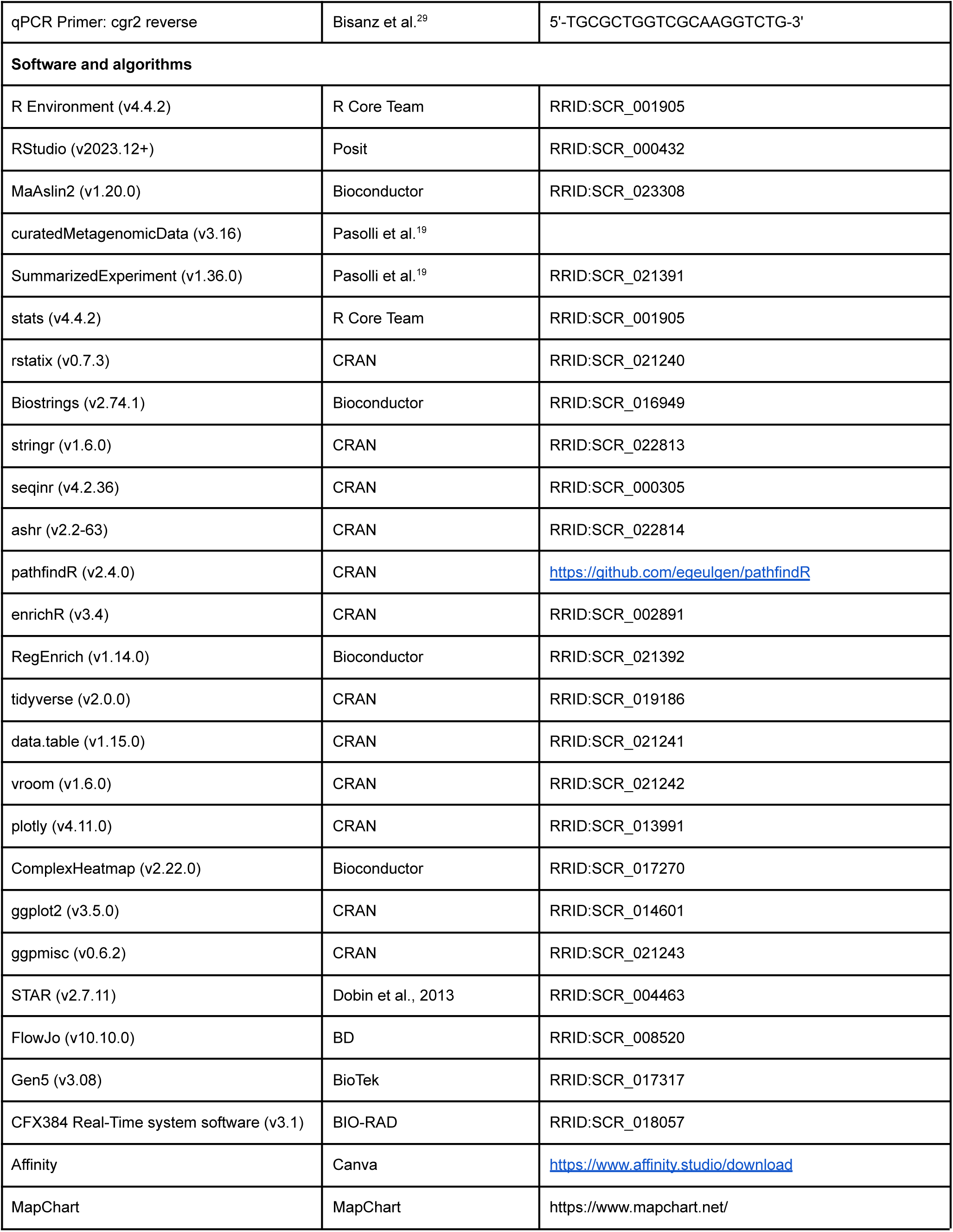
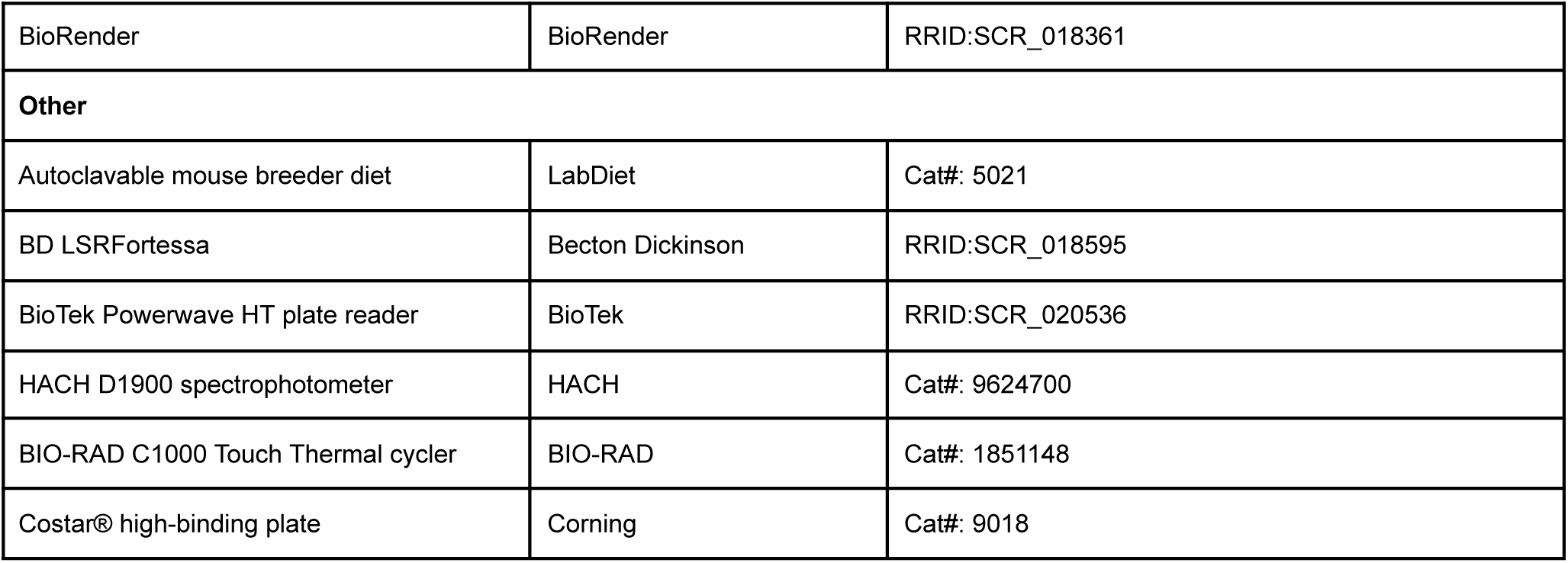

#### EXPERIMENTAL MODEL AND STUDY PARTICIPANT DETAILS

##### Mice

Conventionally raised (CONV-R) C57BL/6J and SJL/J mice were purchased from The Jackson Laboratory and maintained in a reverse light-dark cycle with light phase 7 am–7 pm. All mice were housed at temperatures ranging from 67–74 °F and humidity ranging from 30-70% and fed autoclavable mouse breeder diet *ad libitum*. CONV-R mice were maintained in individual ventilated cages (IVCs) with 10–15 air changes per hour. Germ-free mice were born and maintained in the UCSF Gnotobiotics Core Facility (gnotobiotics.ucsf.edu). Gnotobiotic mice used for EAE experiments were maintained in open-top cages in dedicated, sterile isolators. All other gnotobiotic mice were maintained in IVCs with 10–12 air changes per hour. *tlr2^-^*^/-^ mice (source: JAX) were provided by the Scharschmidt lab. F1 mice were bred to establish a colony and genotyped according to Jackson Labs protocol^53^. Mouse age ranged from 6–9 weeks old and for all experiments mice were assigned into groups to achieve a similar age distribution between groups. No mice used in these experiments were previously used for other procedures or experiments. Full mouse experiment metadata for *in vivo* experiments can be found in **Table S5**. For all experiments involving bacterial colonization, at a minimum, samples from each group were pooled and colonization was confirmed via anaerobic culturing and/or qPCR for an *E. lenta* specific marker gene (*elnmrk1*) or, where applicable, an *E. lenta* strain-specific marker (*cgr2*).

##### Metagenomic dataset sample selection and analysis

We obtained processed metagenomic data using v3.16 of the R package curatedMetagenomicData (cMD), a package providing access to the curated Metagenomics Database repository, which provides published metagenomic data processed using a unified analysis pipeline^19^. Data from all 22,588 samples across 90 studies were filtered for read quality and metadata inclusion. Samples were excluded if any of the following applied: antibiotics use at time of sample collection, mean read depth ≤ 20 million base pairs (bp), median length ≤ 80 bp, non-stool body site, or model adjustment metadata unavailable. After

filtering, 4,681 high-quality samples remained, representing 3,979 individuals from 27 studies. A full list of cMD studies included can be found in **Table S1**. glm from R package stats v4.4.2 was used to fit a logistic model for species prevalence while MaAslin2 from package MaAsLin2^54^ v1.20.0 was used to fit a model for species abundance on the central log ratio (CLR)-normalized data. Covariates were modeled as categorical or continuous variables as appropriate for each modelling framework. Parentheses denote categories used, where applicable. Our models adjusted for sex (male or female), age ((≤1), 1–20 (>1–20), 20–40 (>20–40), 40–60 (>40–60), 60–80 (>60–80), and ≥80 (>80)), BMI (underweight (≤18.5), healthy (>18.5–25), overweight (>25–30), or obese (>30)), continent, health (healthy or ≥1 documented disease), median read length (<100 bases, ≥100 bases), sequencing depth (<100 million, ≥100 million reads), and DNA extraction kit (Qiagen, unknown, other). Our abundance model additionally included study as a random effect, eg: CLR-normalized abundance ∼ sex + age category + BMI category + continent + health category + median read length category + sequencing depth category + DNA extraction kit category + (1|study) whereas our logistic regression model was fit using a fixed-effects framework without random effects, e.g., glm(species prevalence ∼ sex + age + BMI + continent + health + median read length + sequencing depth + DNA extraction kit, family=binomial). Random effects were not included in our prevalence model due to the high sparsity of species-level presence–absence data across studies, which can lead to model non-convergence or unstable parameter estimates in mixed-effects logistic regression. We also obtained the metagenomic sequencing relative abundance table and associated patient metadata from iMSMS^21^, which includes 576 people living with MS (PwMS) and 576 house-hold paired controls. Data were CLR-transformed and a simple linear model was fit adjusting for sex and treatment status (CLR-normalized relative abundance ∼ disease + sex + treatment status).

### METHOD DETAILS

#### DNA extraction and quantification from stool

Mouse fecal samples (30–60 mg of pre-weighed stool) were prepared in ZymoBIOMICS™ Lysis Strip Tubes from the 0.1 mm & 0.5 mm Bead Tube Lysis Rack Kit. Tubes were sealed tightly, suspended in ZymoBIOMICS™ Lysis Buffer, fitted into a ZR-96 BashingBead™ Lysis Rack, and homogenized with bead-beating for 5 minutes. Homogenized lysates were then centrifuged for 5 minutes at 3,000 × g. The supernatant was transferred to 2 mL deep-well plates and the DNA was purified using the ZymoBIOMICS™ 96 MagBead DNA Kit according to the manufacturer’s instructions. Samples were assessed by NanoDrop to confirm isolation success and stored at −20 °C. DNA content was quantified via qPCR using validated primers and a freshly generated standard curve, with positive and negative control samples included in each run. *E. lenta*–specific primers targeting the *E. lenta* marker gene *elnmrk1* were used to quantify *E. lenta* and primers for *E. lenta cgr2* were used to detect *cgr2*^29,55^. The *elnmrk1* primer sequences are forward 5′–GTACAACATGCTCCTTGCGG–3′ (positions 1,548,038–1,548,057) and reverse 5′–CGAACAGAGGATCGGGATGG–3′ (positions 1,548,203–1,548,222), producing a 185 bp amplicon. The *cgr2* primers are forward 5′–CGGCGCGCTTTTTCAGCGTT–3′ (positions 2,959,633–2,959,652) and reverse 5′–TGCGCTGGTCGCAAGGTCTG–3′ (positions 2,959,847–2,959,866), producing a 234-bp amplicon. All genome positions are relative to the National Center for Biotechnology Information [NCBI] RefSeq assembly [GCF] #000024265.1 genome.

#### Experimental autoimmune encephalomyelitis (EAE) in C57BL/6J mice

CONV-R C57BL/6J mice were orally gavaged with 200 µL of ≥ 10^8^ CFU/mL *E. lenta* every other day starting 1–1.5 weeks prior to EAE induction, and gnotobiotic mice colonized with 200 µL of ≥ 10^8^ CFU/mL *E. lenta* 1-3 weeks prior to EAE induction. CONV-R mice were separated into pairs for 3–14 days prior to EAE induction. Induction was performed with minor modifications to previously described protocol^22^. Specifically, immediately before disease induction, unmodified MOG_35–55_ (Tocris or Genscript custom synthesis with <10 Endotoxin Units (EU)/mg) was resuspended in sterile 1× PBS. Complete Freund’s Adjuvant (CFA) was prepared by combining heat-killed *Mycobacterium tuberculosis* with incomplete Freund’s adjuvant. The final emulsion, containing 1 mg/mL MOG_35–55_ and 4 mg/mL MTB, was prepared by passing through the mixture through two 10 mL syringes connected by a three-way stopcock. The emulsion was only used once. 100 μL of the emulsion was injected into mice subcutaneously. Immediately following subcutaneous emulsion injection, 500-1000 ng of pertussis toxin in 100 μL 1× PBS was delivered by intraperitoneal (IP) injection. 48 hours following the initial immunization, an additional 100 µL of 5 μg/mL PTX in PBS was injected IP. Additional modifications specific to gnotobiotic animals are as follows: All procedures were performed in a Class II biosafety cabinet that was UV-irradiated, bleach-decontaminated, and operated with the blower engaged to maintain a sterile work zone. Also, in addition to the water-drop test performed at the time of emulsion preparation, emulsion stability was further confirmed in all cases. Specifically, 1) water-dropped emulsion was saved and examined ≥ one hour after disease induction to ensure its stability by visual inspection for dispersion and 2) gross examination at necropsy to identify an intact, white emulsion depot at the injection site of the emulsion.

#### Experimental autoimmune encephalomyelitis (EAE) in SJL/J mice

Mixed-sex SJL/J mice were purchased from The Jackson Laboratory. Mice were gavaged with 200 µL ≥10^8^ CFU/mL *E. lenta* DSM2243 at 9 weeks of age and induced with EAE two weeks later. Mice that developed tail lesions before or after disease induction were excluded, as recommended by standard protocol^56^. To induce EAE, an emulsion of a 100 µg/100 μL of PLP_139–151_ (Genscript, custom synthesis with >95% purity by high-performance liquid chromatography (HPLC) and <10 EU/mg endotoxin, sequence: HSLGKWLGHPDKF [unmodified]) resuspended in sterile 1× PBS and 4 mg/mL complete Freund’s adjuvant was prepared as described in the previous section. 100 μL of the final emulsion was injected into mice via subcutaneous injection. Directly after PLP injection, 200 ng of PTX in 50 μL of PBS was IP injected. An additional 200 ng PTX in 50 μL of PBS was IP injected the following day. Emulsion stability was confirmed as described in the previous section.

#### Experimental autoimmune encephalomyelitis (EAE) scoring

Mice were weighed and scored for disease every day starting at day of immunization. Singly housed mice were provided dome huts as supplemental enrichment. EAE was scored in accordance with the Hooke EAE rubric^57^ (**Table S6A**). After peak disease, mice were scored according to the Hooke rubric for recovery^57^ (**Table S6B**), as applicable. Investigator scoring animals was blinded to groups for all gnotobiotic experiments. Gnotobiotic mice that died during the course of EAE were excluded from analysis; these mice did not exhibit observable EAE phenotypes, consistent with the generally low prevalence and incomplete penetrance of EAE in gnotobiotic animals^6,8^.

#### Splenocyte isolation

Spleen cells were prepared by gently mashing spleen with a syringe plunger before filtering through a 40 μm filter. Cells were pelleted by centrifugation (900 × g, 5 minutes) and red blood cells lysed using 1 mL 1× red blood cell (RBC) lysis buffer prepared from 10× stock diluted in Milli-Q^®^ ultrapure water. Cells were then pelleted again (900 × g, 5 minutes) and subsequently washed in 1× PBS or enriched for the CD4^+^ fraction using Dynabeads untouched mouse CD4 isolation kit with slight modification to kit specifications. Briefly, cells were suspended in isolation buffer (PBS, 5 mM EDTA, and 0.5% BSA) and incubated at 4 °C with Dynabeads antibody mix (Dynabeads Untouched Mouse CD4 Isolation Kit) for 20 minutes. Cells were then washed and pelleted, followed by a 15 minute room temperature incubation with washed CD4 isolation kit Dynabeads. Dynabeads were removed using magnetic separation, and purified cells were plated in freshly warmed complete Roswell Park Memorial Institute (RPMI) medium (C10), consisting of RPMI 1640 plus glutamine supplemented with 10% FBS, 100 U/mL penicillin–streptomycin, sodium pyruvate, 10 mM HEPES, and 1× non-essential amino acids. For MOG_35-55_ antigen mimicry screening, CD4^+^ T cells of TCR_MOG_ 2D2 mice were isolated from spleen and lymph nodes using a similar protocol and the Miltenyi Biotec CD4 T cell Isolation Kit (Miltenyi Biotec: Cat# 130-104-454).

#### Lamina propria lymphocyte isolation

Gut lamina propria lymphocytes were isolated using only slight modifications of previously described techniques^18,25^. Briefly, the distal third of small intestines (ilea) and colons were splayed longitudinally with adipose tissue and mucus layer removed. During collection, tissues were stored in ice-cold C10. Stored tissues were immediately filtered through a 100 μm filter. Tissue remaining post-filtration was resuspended in 1× Hank’s Balanced Salt Solution containing 5 mM ethylenediaminetetraacetic acid and 1 mM DL-Dithiothreitol before incubating for 45 minutes at 37 °C on a shaker (200 rpm). Each sample supernatant, containing remaining tissue, was filtered through a 100 μm filter into a solution containing 1× Hank’s Balanced Saline Solution with 5% (v/v) fetal bovine serum and a digestion mixture of freshly thawed: 1 U/mL Dispase II, 0.5 mg/mL Collagenase VIII, and 20 μg/mL DNaseI. This solution was then incubated for either 35 minutes (ileum) or 45 minutes (colon) at 37 °C on a shaker set to 200 rpm. The vortexed solution was filtered over a 40 μm cell strainer into 1× PBS. 10% 10× PBS was added to percoll stock to prepare a working solution. Cells were subsequently resuspended in 10 mL of 40% percoll (60% C10, 40% percoll working solution). Carefully, 1 mL 80% percoll (80% percoll working solution, 20% digestion solution) was layered underneath the 40% percoll. Cells were spun at 900 × g for 20 min at 4 °C with no brake and low acceleration. Cells at the interface were collected, washed in 1× PBS and prepared for flow cytometry analysis as described in the following section.

#### Brain lymphocyte isolation

Brain lymphocytes were isolated using only slight modifications of previously described techniques^18^. Briefly, brains were dissected out of the mice and transferred into 4 mL ice cold 1× PBS in 6-well plates. Using scissors or a scalpel, brains were diced into pieces until small enough to homogenize each suspension by passing it through an 18-gauge needle. Each tissue homogenate was then passed through a separate 70 μm filter before filters were washed using 10 mL ice cold 1× PBS. Filtered homogenate was centrifuged at 900 × g for 10 min and supernatant discarded. Cells were resuspended in a 30% percoll mixture prepared using a 30% percoll working solution prepared from percoll stock, 67% heat-inactivated FBS, and 3% 1× PBS. Resuspended cells were overlaid on top of a 70% percoll mixture (70% percoll stock and 30% C10) at room temperature. Cells were spun at 900 × g for 25 minutes at room temperature with no brake and low acceleration. Cells at the interface were collected, washed in 1× PBS and prepared for flow cytometry analysis as described in the following section.

##### Flow cytometry

Lymphocytes were isolated from ileal and colonic lamina propria as described in the previous sections. Cells were stimulated using Cell stimulation cocktail, with protein export blocked using Golgi Plug, for 4 hours at 37 °C. Following stimulation, cells were washed twice with Fluorescence-Activated Cell Sorting (FACS) buffer (1× HBSS, 10 mM HEPES [4-(2-hydroxyethyl)-1-piperazineethanesulfonic acid], 2 mM EDTA, 0.5% heat-inactivated FBS, and 0.02% sodium azide). Cells were stained using the flow protocols detailed in the ADDITIONAL RESOURCES section. For full details on specific methods used and stains by experiment, see **Table S7**.

For basic staining, extracellular antibody cocktails were added to experimental wells in FACS buffer and incubated 20–30 minutes at 4 °C in the dark. Cells were then washed twice, before fixation and permeabilization using a Fixation/Permeabilization Solution Kit. After fixation/permeabilization for 20 minutes at 4 °C, cells were washed and maintained at 4 °C overnight in foil prior to intracellular staining. Intracellular staining was performed using diluted perm/wash buffer as described for extracellular staining.

For transcription factor–optimized staining, cells were resuspended in FACS buffer and incubated with viability and extracellular antibodies for 30 minutes at 4 °C in the dark, followed by washing. Cells were then fixed and permeabilized using the eBioscience Foxp3/Transcription Factor Staining Buffer Set (ThermoFisher, Cat# 00-5523-00) and incubated overnight at 2–8 °C in the dark. All centrifugation steps were performed at 500 × g for 5 minutes at 4 °C, and supernatants were removed using standard aspiration methods for 96-well plates. This workflow encompasses both Transcription Factor Optimized Protocols v1 and v2, with minor variations in incubation time and washing.

Gates were established using freshly prepared isotype and single stain controls, at a minimum. Gating strategies and their correspondence with paper figures are outlined in **Figure S12**. Samples with low viability (ileal and colonic lamina propria: <20% live cells within the singlet cells gate or <10,000 live cells total; brain: <500 live cells total) were excluded from analysis. Summary statistics for live cell counts of included samples by experiment and tissue are provided in **Table S8**. All samples were analyzed on an LSR Fortessa and analyzed using FlowJo v10.10.0.

#### RNA sequencing data analysis

RNA sequencing data were obtained from Alexander et al 2021^25^. Reads were realigned to the *Mus musculus* (house mouse) reference genome GRCm38 (mm10) from the Genome Reference Consortium (GCA_000001635.2; GCF_000001635.20) using industry-standard alignment software (STAR) and Ensembl release 114 (GENCODE M38) gene annotations. Data were analyzed in R (v4.2.2) using DESeq2^58^ v1.46 for differential expression analysis, enrichR^59^ v3.4 for exploratory gene set enrichment analysis, and pathfindR^60^ v2.6.0 for context informed, network-based pathway enrichment analysis. Data visualization was performed using variance-stabilizing transformation (VST)-normalized values, whenever applicable. Core mouse Th signature genes were identified from previously analyzed data sets^61,62^.

#### Strains and bacterial culturing

Bacterial strains (STAR Methods) were cultured at 37 °C in an anaerobic chamber (Coy Laboratory Products; 2%–5% H_2_, 20% CO_2_, balance N_2_). A contamination control was included for all liquid cultures. A chemically-defined media (*Eggerthella lenta* defined media 1, or EDM1, plus 1.25 g/L sodium formate [SIAL: Cat# 71539-500G)^23^ was used for all mouse experiments. Wild-type *E. lenta* DSM2243 with a control plasmid and Δ*cgr* strains were generated as previously described^24^. To prepare heat-killed bacteria and bacterial supernatants for *in vitro* experiments, 5 mL 48-hour stationary phase cultures were prepared. A minimal media was used except for the multi-strain skewing experiment, where a strain-appropriate rich media was used (BHI with 1% L-arginine for DSM2243, DSM11767, DSM110905, DSM110906, DSM16107, and DSM110924; BHI with 0.05% L-cysteine-HCl and 0.0001% weight/volume (w/v) resazurin for DSM120761, and BHI with 0.05% w/v L-cysteine-HCl, 5 μg/mL hemin , and 1 μg/mL menadione for DSM3979. Cultures were pelleted at 2000 rpm for 10 minutes, and then resuspended in 1× PBS. Within each strain, cultures were pooled into a single Eppendorf tube. Tubes were held submerged under water heated to 65 °C on a hotplate, as measured using a calibrated thermometer. The equivalent of 400 µl original overnight culture was immediately removed from each Eppendorf tube for plating and anaerobic incubation at 37 °C for one week. Successful heat-killing of *E. lenta* was validated by confirming <10^3^ CFU/mL. Protein content was normalized to a standard curve made from at least two replicate runs of standards using the BCA Protein assay kit.

#### IFN-γ ELISA

Helper T cells were isolated from healthy mouse spleen as previously described (see “Splenocytes Isolation”) using the Dynabeads^TM^ Untouched^TM^ Mouse CD4 Cell Isolation Kit . Isolated T cells were plated onto a 96-well round-bottom tissue culture plate pre-coated with anti-CD3ε (5 μg/mL) overnight at room temperature. For experiments including multiple animals, cell count was normalized to the lowest viable T cell yield obtained among the mice in that experiment. Cells were provided CD28 (5 μg/mL), for maintenance and incubated under Th1-differentiation conditions consisting of IL-12 (5 ng/mL), IL-2 (2.5 units [U]/mL), and anti-IL-4 (adapted from Huh et al., 2011^63^; concentrations modified after titration). For helper T cell monoculture, T cells purified using the Dynabeads CD4 isolation kit were incubated at 37 °C for 1–2 hours before addition of a standardized quantity of heat-killed bacteria in 1× PBS (Gibco, Cat# 10010-023), TLR2/6 agonist in 1× PBS, or 1× PBS without bacteria. Lot numbers from all PBS used for dilutions were searched online for certificate of analysis (COA) prior to use and endotoxin levels confirmed to be <0.01 EU/mL. No-coat controls, media-only controls, and negative controls were included for every assay. Cells were incubated for 4 days at 37 °C and re-stimulated with Cell Stimulation Cocktail for 16 hours. Subsequently, supernatants were harvested and live cells counted using a hemocytometer. Supernatants were added to a capture-antibody-treated, BSA-blocked Costar® high-binding plate, prepared according to the R&D Systems Mouse IFN-γ DuoSet ELISA Kit protocol. The DuoSet ELISA protocol was followed to prepare plates for IFN-γ detection. To quantify IFN-γ, absorbance was assessed immediately at 570 nm and values assigned based on the average of two standard curves.

#### MOG_35-55_ antigen mimicry screening

We identified putative mimicry epitopes in the *E. lenta* genome (RefSeq assembly [GCF] #000024265.1) using a regex pattern matching search in R. Specifically, we required the antigen to constain a sequence sharing amino acids 2, 5, 7, and 8 with MOG_40-48_ in identical locations^64^. Our criteria for a top hit was: containing sequence with >50% sequence similarity to MOG_40-48_ or previous documented effect in the context of EAE^65^.

For in *vitro coculture*, as previously described^66,67^, CD4^+^ T cells (2 x 10^6^ mL^-1^) were activated with soluble anti–CD3 in the presence of wild–type splenocytes (irradiated at 30 Gy) isolated from TCR^MOG^ 2D2^-^littermates (1 x 10^6^ mL^-1^), which served as antigen–presenting cells to prime the TCR^MOG^ 2D2 CD4^+^ T cells. Peptide or *E. lenta* component in PBS or media or supernatant was added to cells in 1× PBS. Cells were co–cultured for 5 days in lymphocyte culture medium consisting of Dulbecco’s Modified Eagle Medium supplemented with 10% FBS, 50 μM 2–β–mercaptoethanol, vitamin mix solution, 14 μM folic acid solution , 0.7 mM L-arginine, 0.3 mM asparagine mix, 100× non–essential amino acids, penicillin–streptomycin solution, 200 mM L–glutamine, and sodium pyruvate in 6–well plates. Brefeldin A was added during the last 4 h of incubation.

For in *vitro coculture*, as previously described^66^, CD4^+^ T cells (2 x 10^6^ mL^-1^) were activated with soluble anti–CD3_ε_. Cells were stained with viability dye, Fc blocker, and antibodies for surface markers: BV650–anti–CD45.1, PerCP/Cy5.5–anti–CD4, FITC–anti–TCR Vβ11, and, depending on the panel, APC/Cy7–anti–CD25, APC–anti–CD44, PE–anti–CD62L, BV421–anti–CD69, PE–anti–FoxP3, PE/Cy7–anti–Helios, PE–anti–IL–10, APC–anti–IL–17A, APC–anti-RORγT, and eFluor450–anti–IFNγ. For intracellular cytokine and transcription factor staining, the Foxp3 / Transcription Factor Staining Buffer Set and the Fixation/Permeabilization Solution Kit were used.

### QUANTIFICATION AND STATISTICAL ANALYSIS

Statistical tests, the software used, the number of replicates, assumptions made and analysis exclusion criteria are specified in the figure legends or on the plots themselves where possible. FlowJo v10.10.0 was used for gating flow populations as well as generating population scatterplots and histograms. All tests were performed two-tailed, where applicable. Statistical analyses were performed using GraphPad Prism v10.61 and R v4.4.2. All pairwise comparisons between groups were predetermined and performed using rstatix v0.7.2^68^. Unpaired Wilcoxon rank sum tests were used for pairwise comparisons, except when explicitly stated. Longitudinal data was analyzed in GraphPad Prism using a repeated measures mixed model. The response variable was EAE disease score. Fixed effects included experimental group and time (day), with sex included as an additional fixed effect where indicated, and interaction terms were modeled as appropriate. The model accounted for correlations between repeated measurements using mouse as a random effect without assuming sphericity (hence, the Geisser-Greenhouse approximation was not applied). The model used a compound symmetry covariance structure, estimating parameters using Restricted Maximum Likelihood (REML). For correlations, polynomial regression was fitted using ggpmisc v0.6.2^69^. Where applicable, false discovery rate (FDR) correction for multiple hypothesis testing was performed using the Benjamini-Hochberg method, as is noted in figure legends.

### ADDITIONAL RESOURCES

Flow cytometry staining and other immunology protocols are openly available and can be accessed using the links provided below.

● Basic flow staining protocol: https://www.protocols.io/private/F398A866C73D11F0B0220A58A9FEAC02
● Transcription factor optimized flow protocol v1: https://www.protocols.io/private/C1872B30C99411F094BA0A58A9FEAC02
● Transcription factor optimized flow protocol v2): https://www.protocols.io/private/7BC79561C74D11F091A60A58A9FEAC02
● Th1 ELISA: https://www.protocols.io/edit/helper-t-cell-in-vitro-th1-elisa-hfy4b3pyx
● Leukocyte collection and isolation: https://www.protocols.io/edit/leukocyte-isolation-and-tissue-collection-hfyxb3pxp

## Notes

https://www.protocols.io/private/F398A866C73D11F0B0220A58A9FEAC02

https://www.protocols.io/private/C1872B30C99411F094BA0A58A9FEAC02

https://www.protocols.io/private/7BC79561C74D11F091A60A58A9FEAC02

https://www.protocols.io/edit/helper-t-cell-in-vitro-th1-elisa-hfy4b3pyx

https://www.protocols.io/edit/leukocyte-isolation-and-tissue-collection-hfyxb3pxp

### Summary of Updates

This version of the manuscript has been revised to update the low resolution of image files processed in the first iteration of manuscript upload. Please use sepreate high resolution image files for main and supplemental images; thank you!

